# An Atlas of Gene Regulatory Elements in Adult Mouse Cerebrum

**DOI:** 10.1101/2020.05.10.087585

**Authors:** Yang Eric Li, Sebastian Preissl, Xiaomeng Hou, Ziyang Zhang, Kai Zhang, Rongxin Fang, Yunjiang Qiu, Olivier Poirion, Bin Li, Hanqing Liu, Xinxin Wang, Jee Yun Han, Jacinta Lucero, Yiming Yan, Samantha Kuan, David Gorkin, Michael Nunn, Eran A. Mukamel, M. Margarita Behrens, Joseph Ecker, Bing Ren

**Affiliations:** Ludwig Institute for Cancer Research, 9500 Gilman Drive, La Jolla, CA 92093, USA; Center for Epigenomics, Department of Cellular and Molecular Medicine, University of California, San Diego, School of Medicine, La Jolla, CA, USA; Genomic Analysis Laboratory, The Salk Institute for Biological Studies, La Jolla, CA, 92037, USA; Computational Neurobiology Laboratory, Salk Institute for Biological Studies, La Jolla, CA 92037, USA; Department of Cognitive Science, University of California, San Diego, La Jolla, CA 92037, USA; Howard Hughes Medical Institute, The Salk Institute for Biological Studies, La Jolla, CA, 92037, USA; Institute of Genomic Medicine, Moores Cancer Center, University of California San Diego, School of Medicine, La Jolla, CA, USA

**Author notes:** these authors contributed equally (Yang Eric Li, Sebastian Preissl). Correspondence: Bing Ren.

## Abstract

The mammalian cerebrum performs high level sensory, motor control and cognitive functions through highly specialized cortical networks and subcortical nuclei. Recent surveys of mouse and human brains with single cell transcriptomics^1–3^ and high-throughput imaging technologies^4,5^ have uncovered hundreds of neuronal cell types and a variety of non-neuronal cell types distributed in different brain regions, but the cell-type-specific transcriptional regulatory programs responsible for the unique identity and function of each brain cell type have yet to be elucidated. Here, we probe the accessible chromatin in >800,000 individual nuclei from 45 regions spanning the adult mouse isocortex, olfactory bulb, hippocampus and cerebral nuclei, and use the resulting data to define 491,818 candidate *cis* regulatory DNA elements in 160 distinct sub-types. We link a significant fraction of them to putative target genes expressed in diverse cerebral cell types and uncover transcriptional regulators involved in a broad spectrum of molecular and cellular pathways in different neuronal and glial cell populations. Our results provide a foundation for comprehensive analysis of gene regulatory programs of the mammalian brain and assist in the interpretation of non-coding risk variants associated with various neurological disease and traits in humans. To facilitate the dissemination of information, we have set up a web portal (http://catlas.org/mousebrain).

## INTRODUCTION

In mammals, the cerebrum is the largest part of the brain and carries out essential functions such as sensory processing, motor control, emotion, and cognition^6^. It is divided into two hemispheres, each consisting of the cerebral cortex and various cerebral nuclei. The cerebral cortex is further divided into isocortex and allocortex. Isocortex, characterized by six cortical layers, is a phylogenetically more recent structure that has further expanded greatly in primates. It is responsible for sensory motor integration, decision making, volitional motor command and reasoning. The allocortex, by contrast, is phylogenetically the older structure that features three or four cortical layers. It includes the olfactory bulb responsible for processing the sense of smell and the hippocampus involved in learning, memory and spatial navigation.

The cerebral cortex and basal ganglia are made up of a vast number of neurons and glial cells. The neurons can be classified into different types of excitatory projection neurons and inhibitory interneurons, defined by the neural transmitters they produce and their connective patterns with other neurons^7–9^. Understanding how the identity and function of each brain cell type is established during development and modified by experience is one of the fundamental challenges in brain research. Recent single cell RNA-seq and high throughput imaging experiments have produced detailed cell atlases for both mouse and human brains^3–5^,^10–15^, leading to a comprehensive view of gene expression patterns in different brain regions, cell types and physiological states^16–18^. Despite these advances, the gene regulatory programs in most brain cell types have remained to be characterized. A major barrier to the understanding of cell-type specific transcriptional control is the lack of comprehensive maps of the regulatory elements in diverse brain cell types.

Transcriptional regulatory elements recruit transcription factors to exert control of target gene expression in *cis* in a cell-type dependent manner^19^. The regulatory activity of these elements is accompanied by open chromatin, specific histone modifications and DNA hypomethylation^19^. Exploiting these structural features, candidate *cis* regulatory elements (cCREs) have been mapped with the use of tools such as DNase-seq, ATAC-seq, ChIP-seq and Whole genome bisulfite sequencing^20,21^. Conventional assays, typically performed using bulk tissue samples, unfortunately fail to resolve the cCREs in individual cell types comprising the extremely heterogeneous brain tissues. To overcome this limitation, single cell genomic technologies, such as single cell ATAC-seq, have been developed to enable analysis of open chromatin in individual cells^22–28^. These tools have been used to probe transcriptional regulatory elements in the prefrontal cortex^28,29^, cerebellum^29^, hippocampus^30^, forebrain^31^ or the whole brain^24,29^, leading to identification of cell-type specific transcriptional regulatory sequences in these brain regions. These initial studies provided proof of principle for the use of single cell chromatin accessibility assays to resolve cell types and cell-type specific regulatory sequences in complex brain tissues, but the number of cells analyzed, and the *cis* regulatory elements identified so far are still limited.

In the present study, as part of the BRAIN Initiative Cell Census Network, we conducted the most comprehensive analysis to date to identify candidate *cis* regulatory elements (cCRE) in the mammalian brain at single cell resolution. Using a semi-automated single nucleus ATAC-seq (snATAC-seq) procedure^22,31^, we mapped accessible chromatin in >800,000 cells from the mouse isocortex, hippocampus, olfactory bulb, and cerebral nuclei (including striatum and pallidum). We defined 160 sub-types based on the chromatin landscapes and matched 155 of them to previous cell taxonomy of the mouse brain^1^. We delineated the cell-type specificity for >490,000 cCREs that make up nearly 14.8% of the mouse genome. We also integrated the chromatin accessibility data with available brain single cell RNA-seq data to assess their potential role in cell-type specific gene expression patterns, and gain mechanistic insights into the gene regulatory programs of different brain cell types. We further demonstrated that the human counterparts of the identified mouse brain cCREs are enriched for risk variants associated with neurological disease traits in a cell-type-specific and region-specific manner.

## RESULTS

### Single cell analysis of chromatin accessibility of the adult mouse brain

We performed snATAC-seq, also known as sci-ATAC-seq^22,31^, for 45 brain regions dissected from isocortex, olfactory bulb (OLF), hippocampus (HIP) and cerebral nuclei (CNU) (Fig. 1a, Extended Data Figure 1, Supplementary Table 1, see **Methods**) in 8-week-old male mice. Each dissection was made from 600 μm thick coronal brain slices according to the Allen Brain Reference Atlas (Extended Data Figure 1)^32^. For each region, snATAC-seq libraries from two independent biological replicates were generated with a protocol^31^ that had been optimized for automation (Fig. 1a, see **Methods**). The libraries were sequenced, and the reads were deconvoluted based on nucleus-specific barcode combinations. We confirmed that the dataset of each replicate met the quality control metrics (Extended Data Figure 2a-e, see **Methods**). We selected nuclei with at least 1,000 sequenced fragments that displayed high enrichment (>10) in the annotated transcriptional start sites (TSS; Extended Data Figure 2b). We also removed the snATAC-seq profiles likely resulting from potential barcode collision or doublets using a procedure modified from Scrublet^33^ (Extended Data Figure. 2c, see **Methods**). Altogether, we obtained chromatin profiles from 813,799 nuclei with a median of 4,929 fragments per nucleus (Supplementary Table 2). Among them, 381,471 were from isocortex, 123,434 from olfactory area, 147,338 from cerebral nuclei and 161,556 from hippocampus (Fig. 1a, Extended Data Figure 2f). Thus, this dataset represents by far the largest number of chromatin accessibility profiles for these brain areas. Excellent correlation between datasets from similar brain regions (0.92-0.99 for isocortex; 0.89-0.98 for OLF; 0.79-0.98 for CNU; 0.88-0.98 for hippocampus) and between biological replicates (0.98 in median, range from 0.95 to 0.99) indicated high reproducibility and robustness of the experiments (Extended Data Figure 2g).

**Figure 1:**
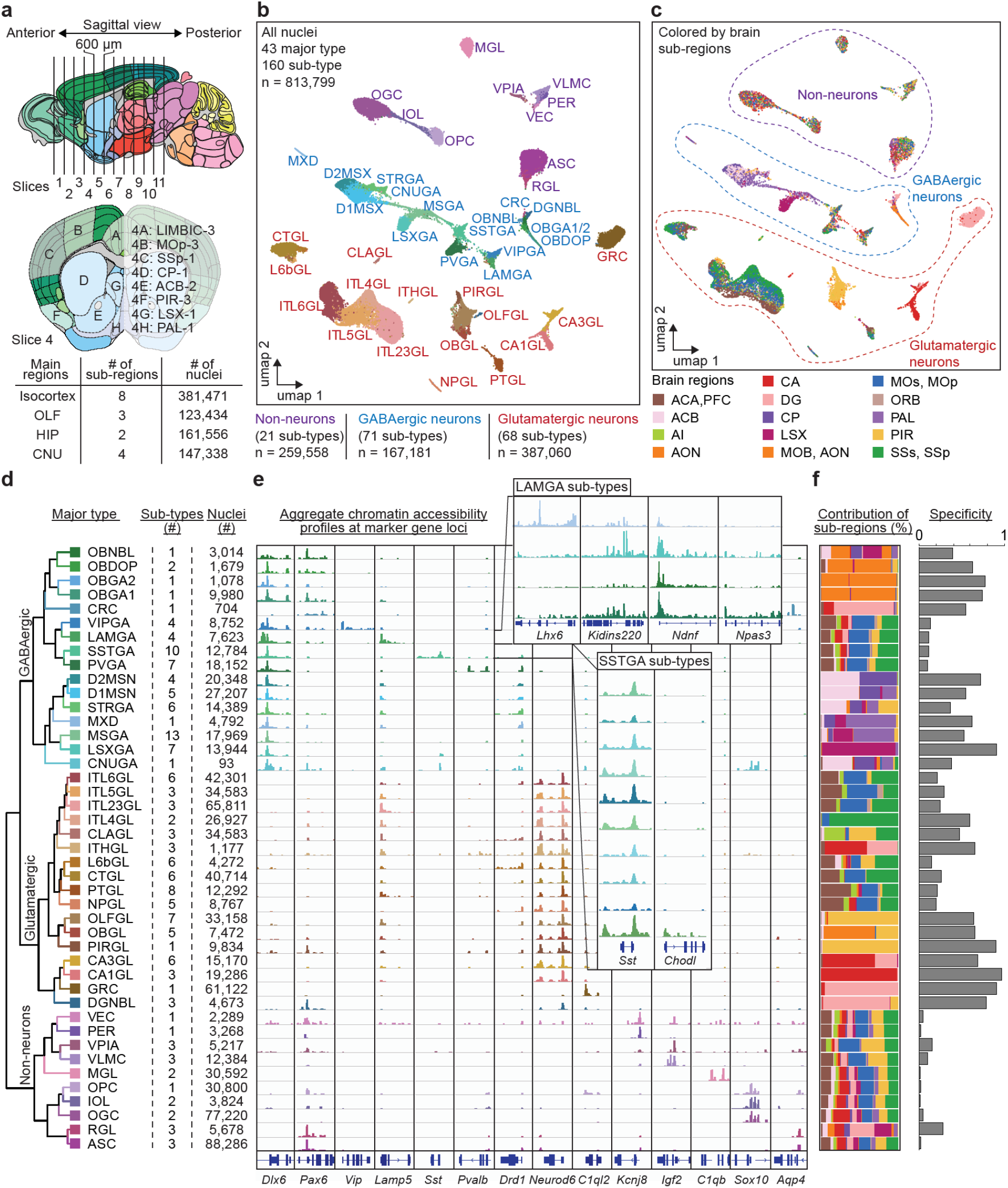
Chromatin accessibility profiling, clustering and annotation of over 800,000 nuclei in adult mouse cerebrum. **a** Schematic of sample dissection strategy. The brain regions studied were dissected from 600 μm-thick coronal slices generated from 8-week-old mouse brains (top panel). A total of 45 regions were dissected according to the Allen Brain Reference Atlas^32^. Shown is the frontal view of slice 4 and the dissected brain regions (middle panel, alphabetically labeled). For example, dissection region 4B: MOp-3 denotes part 3 of the primary motor cortex (MOp) region which corresponds to region B from slice 4. The dissected regions represent 17 sub regions from four main brain areas: isocortex, olfactory bulb (OLF), hippocampus (HIP) and cerebral nuclei (CNU). A detailed list of regions can be found in Supplementary Table 1. **b** Uniform manifold approximation and projection (UMAP)^80^ embedding and clustering analysis of snATAC-seq data from 813,799 nuclei, revealing 43 major types and 160 sub-types assigned to non-neuronal (21, purple), GABAergic (71, blue/green) and Glutamatergic neuron clusters (68, red/brown). Clusters were annotated based on chromatin accessibility at promoter regions and gene bodies of canonical marker genes. Each dot in the UMPA represents a nucleus and the nuclei are colored and labeled by major cluster ID. For example, ITL23GL denotes excitatory neurons from cortex layer 2/3. For a full list and description of cluster labels see Supplementary Table 3. **c** Same embedding as in b but colored by sub-regions, e.g. SSp (primary somatosensory cortex). For a full list of brain regions see Supplementary Table 1. Dotted lines demark major cell classes. **d** Hierarchical organization of cell clusters based on chromatin accessibility depicting level 1 and 2 clusters (left panel). Each major type represents 1-10 sub-types (middle). Total number of nuclei per major type ranged from 93 to 88,286 nuclei (right). For a full list and description of cluster labels see Supplementary Table 2. **e** Genome browser tracks of aggregate chromatin accessibility profiles for each major cell cluster at selected marker gene loci that were used for cell cluster annotation. The inlets highlight the 10 subtypes of *Sst*+ (SSTGA) inhibitory neurons including Chodl-Nos1 neurons (bottom track in SSTGA inlet)^35^ and 4 subtypes of *Lamp5*+ (LAMGA) inhibitory neurons including *Lhx6* positive putative chandelier like cells (top track in LAMGA inlet)^3^. For a full list and description of cluster labels see Supplementary Table 3. **f** Bar chart representing the relative contributions of sub-regions to major clusters. Color code is the same as in b. Based on these relative contributions, an entropy-based specificity score was calculated to indicate if a cluster was restricted to one or a few of the profiled regions (high score) or broadly distributed (low score). Several neuronal types showed high regional specificity whereas non-neuronal types were mostly unspecific. Glutamatergic neurons showed higher regional specificity than GABAergic neurons consistent with transcriptomic analysis^3^.

### Clustering and annotation of mouse brain cells based on open chromatin landscapes

We carried out iterative clustering with the software package SnapATAC^34^ to classify the 813,799 snATAC-seq profiles into distinct cell groups based on the similarity of chromatin accessibility profiles (Fig. 1b-e, Supplementary Table 2 and 3, see **Methods**)^34^. SnapATAC clusters chromatin accessibility profiles using a nonlinear dimensionality reduction method that is highly robust to noise and perturbation^34^. We performed three iterative rounds of clustering, first separating cells into three broad classes, then into major types within each class, and finally into more sub-types. In the first iteration, we grouped cells into glutamatergic neurons (387,060 nuclei, 47.6%), GABAergic neurons (167,181 nuclei, 20.5%) and non-neuronal cells (259,588 nuclei, 31.9%; Fig. 1b-d). For each main cell class, we performed a second round of clustering. We identified a total of 43 major types including distinct layer-specific cortical neurons, hippocampal granular cells (GRC) and striatal D1 and D2 medium spiny neurons (D1MSN, D2MSN; Fig 1b, d) which were annotated based on chromatin accessibility in promoters and gene bodies of known marker genes (Fig. 1 e)^1,3^. Finally, for each major type we conducted another round of clustering to reveal sub-types. For example, *Lamp5^+^* neurons (LAMGA) and *Sst^+^* neurons (SSTGA) were further divided into sub-types (Fig. 1d, e, Supplementary Table 3)^3,35^. One of the LAMGA subtypes showed accessibility at *Lhx6* and therefore might resemble an unusual transcriptomically defined putative chandelier-like cell type with features from caudal ganglionic and medial ganglionic eminence (Fig. 1b, e)^3^. Similarly, using this third layer clustering we found one SSTGA subpopulation with accessibility at *Chodl* locus which resembles long range projecting GABAergic neurons (Fig. 1b, e)^35^. Altogether, we were able to resolve 160 sub-types, with the number of nuclei in each group ranging from 93 to 75,474 and a median number of 5,086 nuclei per cluster (Supplementary Table 3).

We constructed a hierarchical dendrogram to represent the relative similarity in chromatin landscapes among the 43 major cell groups (Fig. 1d, Extended Data Figure 3). This dendrogram captures known organizing principles of mammalian brain cells: Neurons are separated from non-neuronal types followed by separation of neurons based on neurotransmitter types (GABAergic versus glutamatergic) and finally into more specified cell types which might resemble the developmental origins (Fig. 1d)^3^. Consistent with previous reports of brain cell types, we found that non-neuronal cells were broadly distributed in all regions while several classes of glutamatergic neurons and GABAergic neurons showed regional specificity (Fig. 1c, f, Extended Data Figure 4)^3^. We also found that glutamatergic neuron types showed more regional specificity than GABAergic types, consistent with transcriptomic analysis (Fig. 1c, f, Extended Data Figure 4)^3^.

The chromatin-defined cell types matched well with the previously reported taxonomy based on transcriptomes and DNA methylomes^3,36^ (see companion manuscript by Liu, Zhou et al.^37^). To directly compare our single nucleus chromatin-derived cell clusters with the single cell transcriptomics defined taxonomy of the mouse brain^1^, we first used the snATAC-seq data to impute RNA expression levels according to the chromatin accessibility of gene promoter and gene body as described previously (Seurat package^38^). We then performed integrative analysis with scRNA-seq data from matched brain regions of the Mouse Brain Atlas^1^. We found strong correspondence between the two modalities which was evidenced by co-embedding of both transcriptomic (T-type) and chromatin accessibility (A-type) cells in the same joint clusters (Fig. 2a-c, Supplementary Table 4, see **Methods**). For this analysis, we examined GABAergic neurons, glutamatergic neurons and non-neuronal cell classes separately (Fig. 2a-c, Supplementary Table 4, see **Methods**). For 155 of 160 types defined by snATAC-seq (A-Type), we could identify a corresponding cell cluster defined using scRNA-seq data (T-Type, Fig. 2d, e); conversely, for 84 out of 100 T-types we identified one, or in some cases more, corresponding A-types (Fig. 2d, f). Of note, two clusters fell into different classes. The Cajal-Retzius cells (CRC) was part of the GABAergic class in A-type but glutamatergic class in T-type and one small non-neuronal A-type cluster, VPIA3 (Vascular and leptomeningeal like cells) co-clustered with CRC T-type (Fig. 2d). Nevertheless, the general agreement between the open chromatin-based clustering and transcriptomics-based clustering laid the foundation for integrative analysis of cell-type specific gene regulatory programs in the mouse brain using single cell RNA and single nucleus chromatin accessibility assays, as for the mouse primary motor cortex^15^.

**Figure 2:**
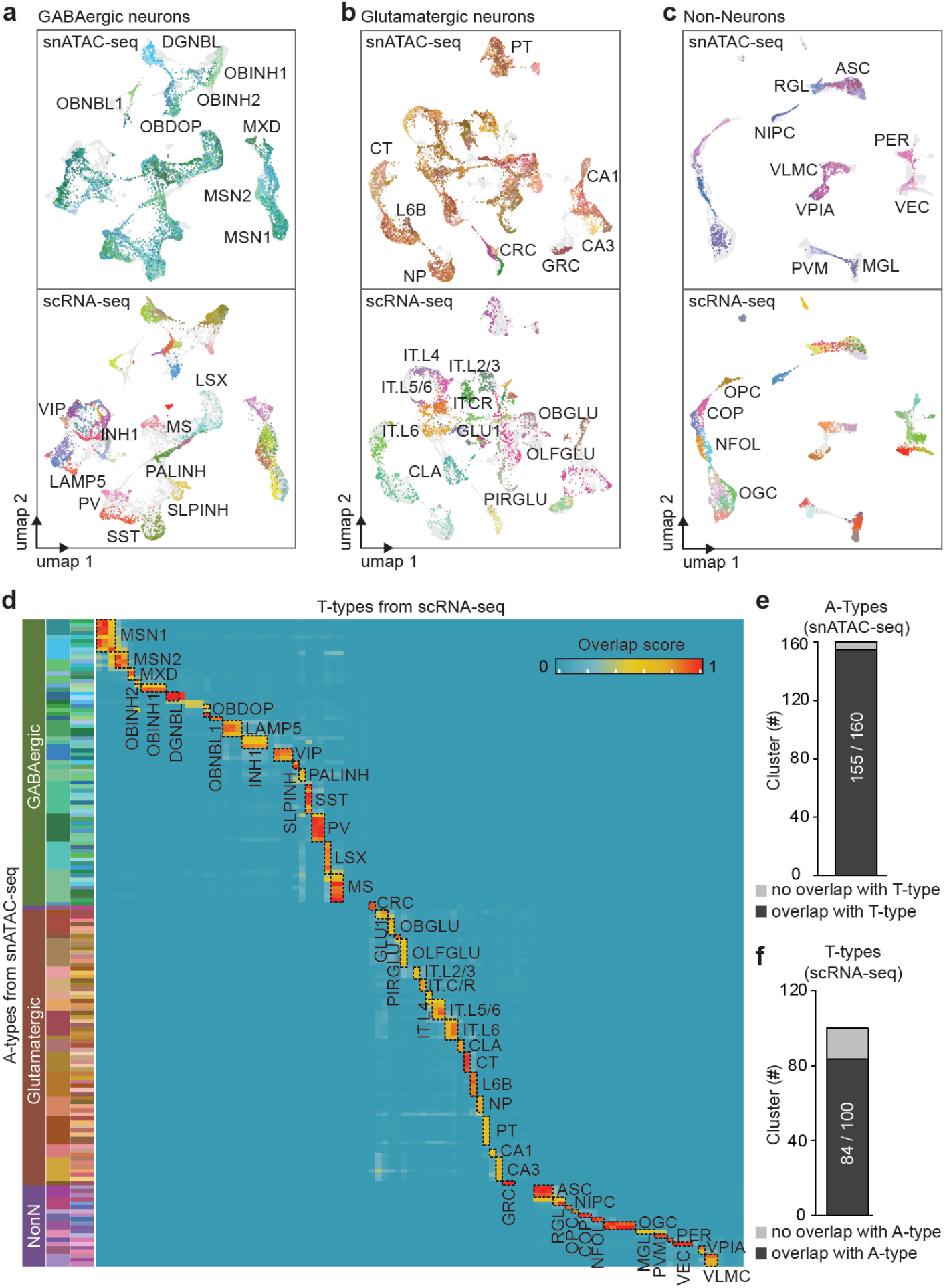
Alignment of chromatin-based cell clustering to scRNA-seq-based cell type taxonomy. **a-c** SnATAC-seq data were integrated with scRNA-seq profiles from matched brain regions^1^ using the Seurat package^38^. Uniform manifold approximation and projections (UMAPs)^80^ illustrate co-embedding of snATAC-seq and scRNA-seq datasets from three main cell classes, namely c GABAergic neurons, **d** glutamatergic neurons, and **e** non-neurons (top: colored by snATAC-seq clusters (A-type), bottom: colored by scRNA-seq clusters (T-type); labelling denotes integrated A/T-types). **d** Heatmap illustrating the overlap between A-type and T-type cell cluster annotations. Each row represents a snATAC-seq sub-type (total of 160 A-types) and each column represents scRNA-seq cluster (total of 100 T-types). The overlap between original clusters and the joint cluster was calculated (overlap score) and plotted on the heatmap. Joint clusters with an overlap score of >0.5 are highlighted using black dashed line and labeled with joint cluster ID. For a full list of cell type labels and description see Supplementary Table 4. **e, f** Bar plots indicating the number of clusters that overlapped (dark grey) and that did not overlap (light grey) with clusters from the other modality. **e** 155 out of 160 A-types had a matching T-type. **f** 84 out of 100 T-types had a matching A-type.

### Identification of cCREs in different mouse brain cell types

To identify the cCREs in each of the 160 A-types defined from chromatin landscapes, we aggregated the snATAC-seq profiles from the nuclei comprising each cell cluster and determined the open chromatin regions with MACS2^39^. We then selected the genomic regions mapped as accessible chromatin in both biological replicates, finding an average of 93,775 (range from 50,977 to 136,962) sites (500-bp in length) in each sub-type. We further selected the elements that were identified as open chromatin in a significant fraction of the cells in each sub-type (FDR >0.01, zero inflated Beta model, see **Methods**), resulting in a union of 491,818 open chromatin regions. These cCREs occupied 14.8% of the mouse genome (Supplementary Table 5 and 6).

96.3% of the mapped cCREs were located at least 1 kbp away from annotated promoter regions of protein-coding and lncRNA genes (Gencode V16) (Fig. 3a)^40^. Several lines of evidence support the function of the identified cCREs. First, they largely overlapped with the DNase hypersensitive sites (DHS) previously mapped in a broad spectrum of bulk mouse tissues and developmental stages by the ENCODE consortium (Fig. 3b)^41,42^. Second, they generally showed higher levels of sequence conservation than random genomic regions with similar GC content (Fig 3c). Third, they were enriched for active chromatin states or potential insulator protein binding sites previously mapped with bulk analysis of mouse brain tissues (Fig. 3d)^43–45^.

**Figure 3:**
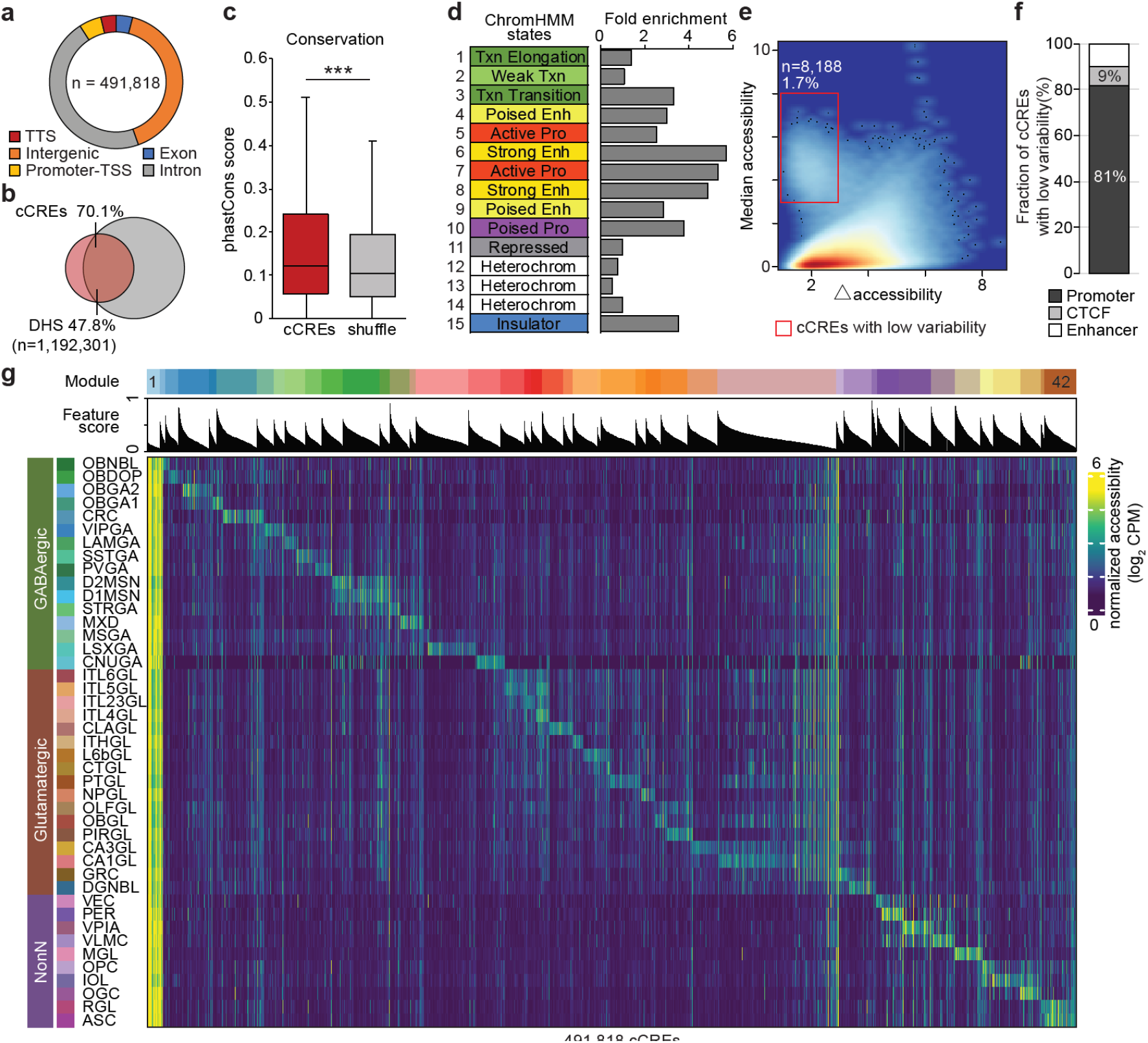
Characterization of candidate *cis* regulatory elements identified in mouse cerebral cell types. **a** Fraction of the identified cCREs that overlap with annotated transcriptional start sites (TSS), introns, exons, transcriptional termination sites (TTS) and intergenic regions in the mouse genome. **b** Venn diagram showing the overlap between cCREs and DNase hypersensitive sites (DHS) from developmental and adult mouse tissue from the SCREEN database^42^. **c** Box-Whisker plot showing that sequence conservation measured by PhastCons score^81^ is higher for cCREs than the controls consisting of GC-matched random genomic sequences (*** p <0.001, Wilcoxon rank sum test, the box is drawn from lower quartile (Q1) to upper quartile (Q3) with a horizontal line drawn in the middle to denote the median, whiskers with maximum 1.5 IQR). d Enrichment analysis of cCREs with a 15-state ChromHMM model^45^ in the mouse brain chromatin^43^. **e** Density map showing two main groups of elements based on the median accessibility and the range of chromatin accessibility variation (maximum – minimum) across cell clusters for each cCRE. Each dot represents a cCRE. Red box highlights elements with low chromatin accessibility variability across clusters. **f** 81 % of sites with low variability (red box in e) overlapped promoters, 10 % enhancers and 9 % CTCF regions. **g** Heatmap showing association of 43 major cell types (rows) with 42 *cis* regulatory modules (top). Each column represents one of 491,818 cCREs. These cCREs were combined into *cis* regulatory modules based on accessibility patterns across major cell types. For each cCRE a feature score was calculated to represent the specificity for a given module. Module 1 comprised invariable elements and was enriched for promoters. For a full list and description of cell cluster labels see Supplementary Table 3, for a full list of cluster-module association see Supplementary Table 7 and for association of cCREs to modules see Supplementary Table 8. CPM: counts per million.

To define the cell-type specificity of the cCREs, we first plotted the median levels of chromatin accessibility against the maximum variation for each element (Fig 3e). We found that the majority of cCREs displayed highly variable levels of chromatin accessibility across the brain cell clusters identified in the current study, with the exception for 8,188 regions that showed accessible chromatin in virtually all cell clusters (Fig 3e). The invariant cCREs were highly enriched for promoters (81%), with the remainder including CTC-binding factor (CTCF) binding sites and strong enhancers (Fig 3f). To more explicitly characterize the cell-type specificity of the cCREs, we used non-negative matrix factorization to group them into 42 modules, with elements in each module sharing similar cell-type specificity profiles. Except for the first module (M1) that included mostly cell-type invariant cCREs, the remaining 41 modules displayed highly cell-type restricted accessibility (Fig. 3g, Supplementary Table 7, 8). These cell-type restricted modules were enriched for transcription factor motifs recognized by known transcriptional regulators for such as the SOX family factors for oligodendrocytes OGC (Supplementary Table 9)^46,47^. We also found strong enrichment for the known olfactory neuron regulator LIM homeobox factor LHX2 in module M5 which was associated with GABAergic neurons in the olfactory bulb (OBGA1) (Supplementary Table 9)^48^.

### Integrative analysis of chromatin accessibility and gene expression across mouse brain cell types

To dissect the transcriptional regulatory programs responsible for cell-type specific gene expression patterns in the mouse cerebrum, we carried out integrative analysis combining the single nucleus ATAC-seq collected in the current study with single cell RNA-seq data from matched brain regions^1^. Enhancers can be linked to putative target genes by measuring co-accessibility between enhancer and promoter regions of putative target genes and co-accessible sites tend to be in physical proximity in the nucleus^49^. Thus, we first identified pairs of co-accessible cCREs in each cell cluster using Cicero^49^ and inferred candidate target promoters for distal cCRE located more than 1 kbp away from annotated transcription start sites in the mouse genome (Fig. 4a, see **Methods**)^40^. We determined a total of 813,638 pairs of cCREs within 500 kbp of each other, and connected 261,204 cCREs to promoters of 12,722 genes (Supplementary Table 10).

**Figure 4:**
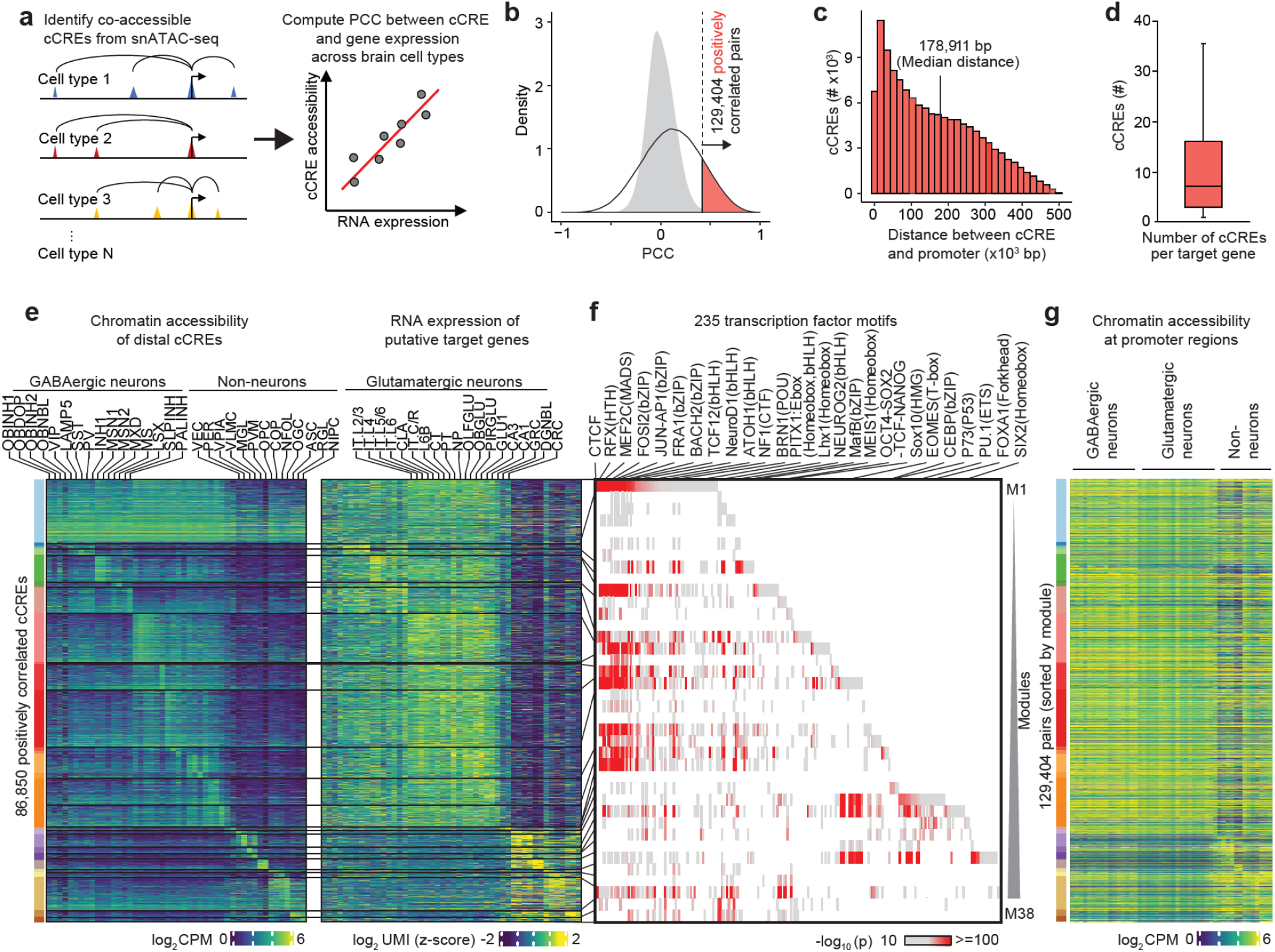
Identification and characterization of putative enhancer-gene pairs. **a** Schematic overview of the computational strategy to identify cCREs that are positively correlated with transcription of target genes. The cCREs were first assigned to putative target gene promoters in specific cell clusters using co-accessibility analysis with Cicero^49^. Next, chromatin accessibility at cCREs was correlated with RNA-seq signals of the putative target gene across different cell clusters (PCC: Pearson correlation coefficient). **b** Detection of putative enhancer-gene pairs. 129,404 pairs of positively correlated cCRE and genes (highlighted in orange) were identified using an empirically defined significance threshold of FDR<0.01 (see **Methods**). Grey filled curve shows distribution of PCC for randomly shuffled cCRE-gene pairs. **c** Histogram illustrating distance between positively correlated distal cCRE and putative target gene promoters. Median distance was 178,911 bp. **d** Box-Whisker plot showing that genes were linked with a median of 7 putative enhancers (box is drawn from Q1 to Q3 with a horizontal line drawn in the middle to denote the median, whiskers with maximum 1.5 IQR). **e** Heatmap of chromatin accessibility of 86,850 putative enhancers across cell clusters (left) and expression of 10,604 linked genes (right). Note genes are displayed for each putative enhancer separately. For association of modules with cell types see Supplementary Table 11 and association of individual putative enhancer with modules see Supplementary Table 13. CPM: counts per million, UMI: unique molecular identifier. **f** Enrichment of known transcription factor motifs in distinct enhancer-gene modules. Displayed are known motifs from HOMER^46^ with enrichment p-value <10^−10^. Motifs were sorted based on module. For full list see Supplementary Table 14. g Accessibility at promoter regions across joint A/T-types, same order as **e**.

Next, we sought to identify the subset of cCREs that might increase expression of putative target genes and therefore function as putative enhancers in neuronal or non-neuronal types. To this end, we first identified distal cCREs for which chromatin accessibility was correlated with transcriptional variation of the linked genes in the joint cell clusters as defined above (Fig. 2a). We computed Pearson correlation coefficients (PCC) between the normalized chromatin accessibility signals at each cCRE and the RNA expression of the predicted target genes across these cell clusters (Fig. 4a, b). As a control, we randomly shuffled the cCREs and the putative target genes and computed the PCC of the shuffled cCRE-gene pairs (Fig. 4b, see **Methods**). This analysis revealed a total of 129,404 pairs of positively correlated cCRE (putative enhancers) and genes at an empirically defined significance threshold of FDR < 0.01 (Supplementary Table 10). These included 86,850 putative enhancers and 10,604 genes (Fig. 4b). The median distances between the putative enhancers and the target promoters was 178,911 bp (Fig. 4c). Each promoter region was assigned to a median of 7 putative enhancers (Fig. 4d), and each putative enhancer was assigned to one gene on average. To investigate how the putative enhancers may direct cell-type specific gene expression, we further classified them into 38 modules, by applying non-negative matrix factorization to the matrix of normalized chromatin accessibility across the above joint cell clusters. The putative enhancers in each module displayed a similar pattern of chromatin accessibility across cell clusters to expression of putative target genes (Fig 4e, Supplementary Table 11 and 13). This analysis revealed a large group of 12,740 putative enhancers linked to 6,373 genes expressed at a higher level in all neuronal cell clusters than in all non-neuronal cell types (module M1, top, Fig. 4e). It also uncovered modules of enhancer-gene pairs that were active in a more restricted manner (modules M2 to M38, Fig 4e). For example, module M33 was associated with perivascular microglia (PVM). Genes linked to putative enhancers in this module were related to immune gene and the putative enhancers were enriched for the binding motif for ETS-factor PU.1, a known master transcriptional regulator of this cell lineage (Fig. 4e, f, Supplementary Table 13 and 14)^50^. Similarly, module M35 was strongly associated with oligodendrocytes (OGC) and the putative enhancers in this module were enriched for motifs recognized by the SOX family of transcription factors (Fig. 4e, f, Supplementary Table 14)^47^. We also identified module M15 associated with several cortical glutamatergic neurons (IT.L2/3,IT.L4,IT.L5/6,IT.L6), in which the putative enhancers were enriched for sequence motifs recognized by the bHLH factors NEUROD1 (Fig. 4e, f, Supplementary Table 14)^51^. Another example was module M10 associated with medium spiny neurons (MSN1 and 2), in which putative enhancers were enriched for motif for the MEIS factors, which play an important role in establishing the striatal inhibitory neurons (Fig. 4e, f, Supplementary Table 14)^52^. Notably and in stark contrast to the striking differences at putative enhancers, the chromatin accessibility at promoter regions showed little variation across cell types (Fig. 4g). This is consistent with the paradigm that cell-type-specific gene expression patterns are largely established by distal enhancer elements^42,53^.

### Distinct groups of transcription factors act at the enhancers and promoters in the pan-neuronal gene module

As shown above, genes associated with module M1 are preferentially expressed in both glutamatergic and GABAergic neurons, but not in glial cell types (Fig. 4e). *De novo* motif enrichment analysis of the 12,740 cCREs or putative enhancers in this module showed dramatic enrichment of sequence motifs recognized by the transcription factors CTCF, RFX, MEF2 (Supplementary Table 15), as well as many known motifs for other transcription factors (Fig. 4f, Fig. 5a, Supplementary Table 14). CTCF is a ubiquitously expressed DNA binding protein with a well-established role in transcriptional insulation and chromatin organization^54^. Recently, it was recognized that CTCF also promotes neurogenesis by binding to promoters and enhancers of proto-cadherin alpha gene cluster and facilitating enhancer-promoter contacts^55,56^. In the current study we found putative enhancers with CTCF motif for 2,601 genes that were broadly expressed in both inhibitory and excitatory neurons (Fig. 4e, 5b), and involved in multiple neural processes including axon guidance, regulation of axonogenesis, and synaptic transmission (Fig. 5c, Supplementary Table 16). For example, we found one CTFC peak overlapping a distal cCRE positively correlated with expression of *Lgi1* which encodes a protein involved in regulation of presynaptic transmission^57^ (Fig 5d). The RFX family of transcription factors are best known to regulate the genes involved in cilium assembly pathways^58^. Unexpectedly, we found the RFX binding sequence motif to be strongly enriched at the putative enhancers for genes encoding proteins that participate in postsynaptic transmission, postsynaptic transmembrane potential, mitochondrion distribution, and receptor localization to synapse (Fig. 5c, Supplementary Table 16). For example, we found RFX motif in a distal cCRE positively correlated with expression of *Kif5a* which encodes a protein essential for GABAA receptor transport (Fig. 5e)^59^. This observation thus suggests a role for RFX family of transcription factors in regulation of synaptic transmission pathways in mammals. Similar to CTCF and RFX, the MEF2 family transcription factors have also been shown to play roles in neurodevelopment and mental disorders^60^. Consistent with this, the genes associated with putative enhancers containing MEF2 binding motifs were selectively enriched for those participating in positive regulation of synaptic transmission, long-term synaptic potentiation, and axonogenesis (Fig. 5c, Supplementary Table 16). For example, we found a distal cCRE harboring a MEF2 motif positively correlated with expression of *Cacng2* which encodes a calcium channel subunit that is involved in regulating gating and trafficking of glutamate receptors (Fig 5f)^61^. Notably, in types with high accessibility levels, cCREs and promoters of putative target genes also showed low levels of DNA methylation (Fig. 5d-f, see companion manuscript by Liu, Zhou et al. 2020^37^).

**Figure 5:**
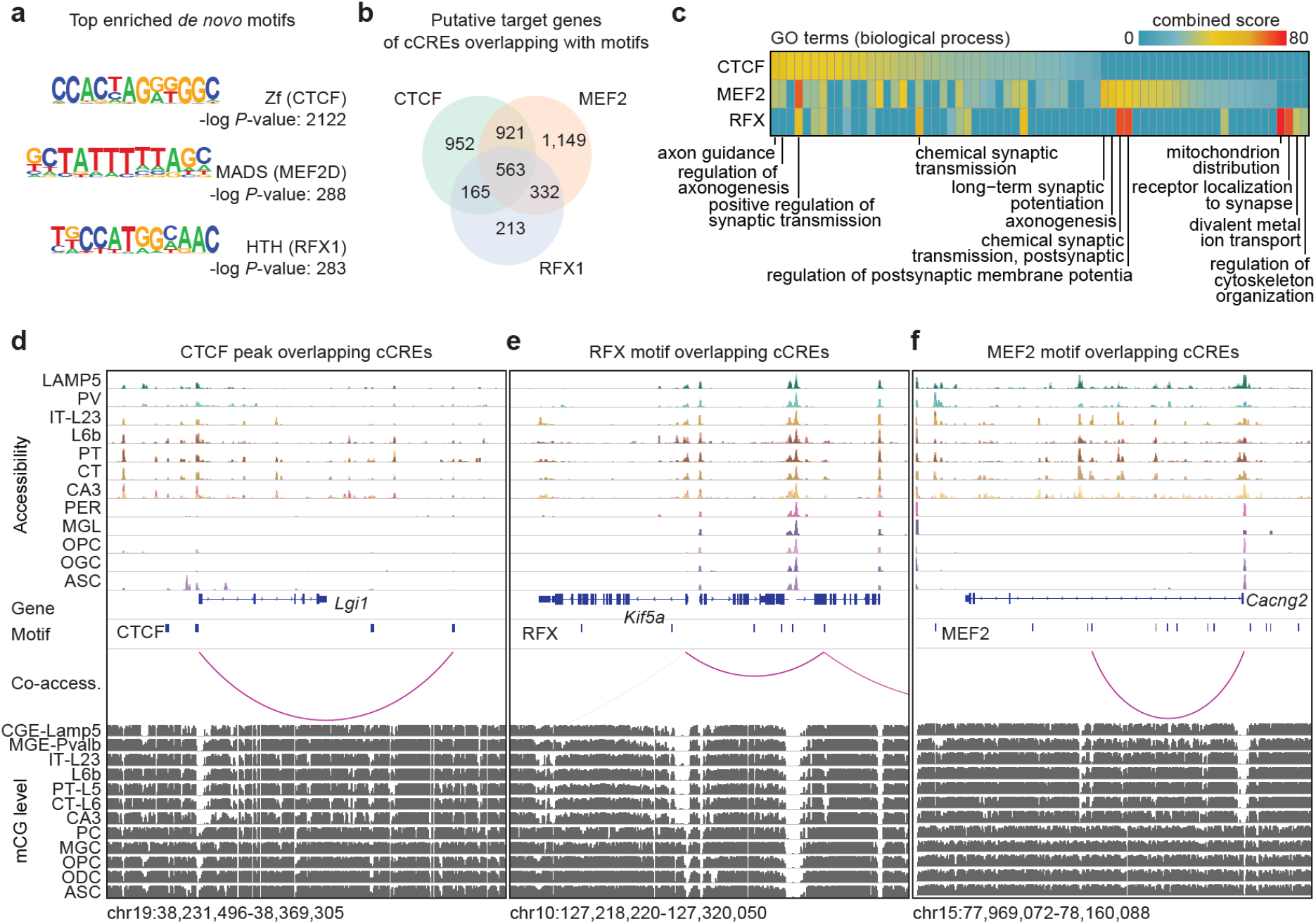
Transcription factors involved in a pan neuronal gene regulatory program. **a** Enrichment of sequence motifs for CTCF, MEF2 and RFX from de novo motif search in the putative enhancers of module M1 using HOMER^46^. For a full list see Supplementary Table 16. **b** Venn diagram illustrating the overlap of putative target genes of cCREs containing binding sites for MEF2, RFX and CTCF, respectively. **c** Gene ontology (GO) analysis of the putative target genes of each factor in module M1 was performed using Enrichr^82^. The combined score is the product of the computed p value using the Fisher exact test and the z-score of the deviation from the expected rank^82^. **d-f** Examples distal cCRE overlapping peaks/motifs and positively correlated putative target genes. For CTCF, cCREs were intersected with peak calls from ChIP-seq experiments in the adult mouse brain^43^ (**d**) and cCREs overlapping RFX (**e**) and MEF2 (**f**) were identified using de novo motif search in HOMER^46^. Genome browser tracks displaying chromatin accessibility, mCG methylation levels (see companion manuscript by Liu, Zhou et al. 2020^37^) and positively correlated cCRE and genes pairs.

Interestingly, motif analysis of promoters of genes linked to cCREs in the module M1 revealed the potential role of very different classes of transcription factors in neuronal gene expression. Among the top ranked transcription factor motifs are those recognized by CREB (cAMP-response elements binding protein), NF-κB, STAT3 and CLOCK transcription factors (Supplementary Table 17). Enrichment of CREB binding motif in module M1 gene promoters is consistent with its well-documented role in synaptic activity-dependent gene regulation and neural plasticity^62,63^. Enrichment of NF-κB^64^, STAT3^65^ and CLOCK^66^ binding motifs in the module M1 gene promoters is interesting, too, as it suggests potential roles for additional extrinsic signaling pathways, i.e. stress, interferon, circadian rhythm, respectively, in the regulation of gene expression in neurons.

### Non-coding variants associated with neurological traits and diseases are enriched in the human orthologs of the mouse brain cCREs in a cell type-specific manner

Genome-wide association studies (GWASs) have identified genetic variants associated with many neurological disease and traits, but interpreting the results have been challenging because most variants are located in non-coding parts of the genome that often lack functional annotations and even when a non-coding regulatory sequence is annotated, its cell-type specificity is often not well known^67,68^. To test if our maps of cCREs in different mouse brain cell type could assist the interpretation of non-coding risk variants of neurological diseases, we identified orthologs of the mouse cCREs in the human genome by performing reciprocal homology search^69^. For this analysis, we found that for 69.2% of the cCREs, human genome sequences with high similarity could be identified (> 50 % of bases lifted over to the human genome, Fig. 6a). Supporting the function of the human orthologs of the mouse brain cCREs, 83.0% of them overlapped with representative DNase hypersensitivity sites (rDHSs) in the human genome^41,42^. Next, we performed linkage disequilibrium (LD) score regression (LDSC)^70^ to determine associations between different brain regions and distinct GWAS traits (Fig. 6b, Extended Data Figure 5). We found a significant enrichment of cCREs from 36 out of 45 brain regions for risk variants of Schizophrenia (Fig. 6b). In fact, most neurological traits showed widespread enrichment across brain regions, but a few like ADHD (Attention deficit hyperactivity disorder) showed some regional enrichment patterns (Fig. 6b).

**Figure 6:**
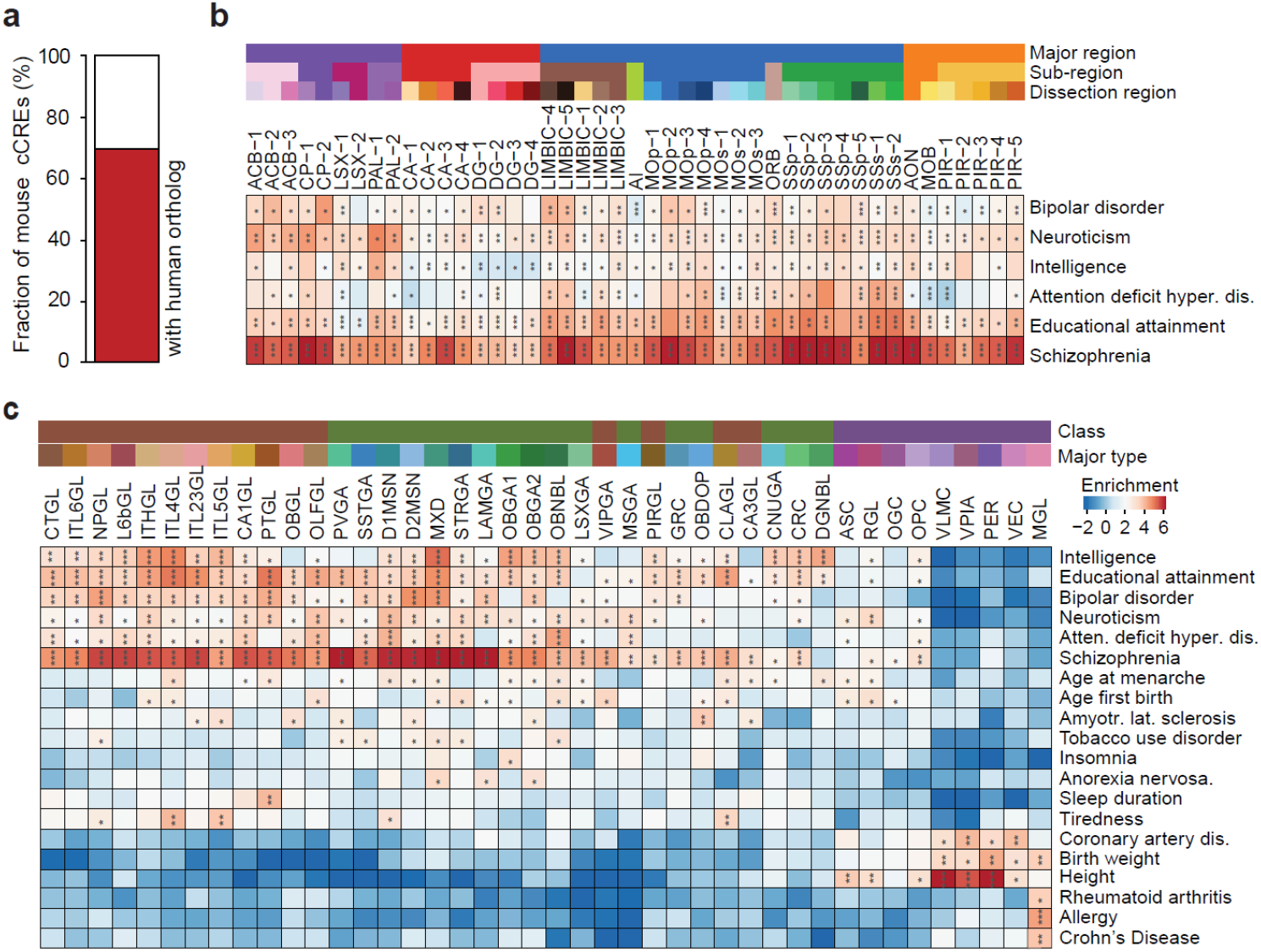
Association of different brain regions and cell types with risk variants for neurological diseases and traits. **a** For 69.2 % of cCREs identified in the current study, we found a human ortholog (> 50 % of bases lifted over to the human genome). **b** Brain-region-specific enrichment of sequence variants associated with indicated neurological traits and diseases (* FDR < 0.05, ** FDR < 0.01, ***FDR < 0.001). Displayed are all regions and all tested phenotypes with at least one significant association. **c** Enrichment of sequence variants associated with the indicated traits/disease in the human orthologs of cCREs in major mouse cerebral cell types (* FDR < 0.05, ** FDR < 0.01, ***FDR < 0.001). Displayed are all major cell clusters and tested traits/diseases with at least one significant association (FDR < 0.05). A detailed list of regions can be found in Supplementary Table 1 and a full list of cell cluster labels can be found in Supplementary Table 3.

We also performed LDSC analysis and found significant associations between 20 neuronal and non-neuronal traits and cCREs found in one or more major cell types (Fig. 6c). We observed widespread and strong enrichment of genetic variants linked to psychiatric and cognitive traits such as major depressive disorder, intelligence, neuroticism, educational attainment, bipolar disorder and schizophrenia in cCREs across various neuronal cell types (Fig. 6c). Other neurological traits, such as attention deficit hyperactivity disorder, chronotype, autism spectrum disorder and insomnia were associated with specific neuronal cell-types in cerebral nuclei and hippocampus (Fig. 6c). Schizophrenia risk variants were not only enriched in cCREs in all excitatory neurons, but also in certain inhibitory neuron sub-types (Fig. 6c)^71^. The strongest enrichment of heritability for bipolar disorder was in elements mapped in excitatory neurons from isocortex (Fig. 6c). Risk variants of tobacco use disorder showed significant enrichment in the cell types from striatum, a cerebral nucleus previously implicated in addiction (Fig. 6c)^72^. Interestingly, cCREs of non-neuronal mesenchymal cells were not enriched for neurological traits but showed enrichment for cardiovascular traits such as coronary artery disease (Fig. 6c). Similarly, variants associated with height were also significant in these cell types (Fig. 6c). cCREs in microglia were significantly enriched for variants related to immunological traits like inflammatory bowel disease, Crohn’s disease and multiple sclerosis (Fig. 6c). Notably, most of these patterns were not apparent in the peaks called on aggregated bulk profiles from brain regions (Fig. 6b, Extended Data Fig. 5), demonstrating the value of cell type resolved open chromatin maps which was also highlighted by a recent study using single cell ATAC-seq profiling of human brain which focusing on Alzheimers’ and Parkinson’s disease^73^.

## DISCUSSION

Understanding the cellular and molecular genetic basis of brain circuit operations is one of the grand challenges in the 21^st^ century^12,74^. In-depth knowledge of the transcriptional regulatory program in brain cells would not only improve our understanding of the molecular inner workings of neurons and non-neuronal cells, but could also shed new light into the pathogenesis of a spectrum of neuropsychiatric diseases^75^. In the current study, we report comprehensive profiling of chromatin accessibility at single cell resolution in the mouse cerebrum. The chromatin accessibility maps of 491,818 cCREs, probed in 813,799 nuclei and 160 sub-types representing multiple cerebral cortical areas and subcortical structures, are the largest of its kind so far. The cell type annotation based on open chromatin landscape showed strong alignment with those defined based on single cell transcriptomics^1^, which allowed us to jointly analyse the two molecular modalities across major cell types in the brain and identify putative enhancers for over 10,604 genes expressed in the mouse cerebrum. We further characterized the cell-type-specific activities of putative enhancers, inferred their potential target genes, and predicted transcription factors that act through these candidate enhancers to regulate specific gene modules and molecular pathways.

We identified one large group of putative enhancers for genes that are broadly expressed in GABAergic and glutamatergic neurons, but at low levels or are silenced in all glial cell types. A significant fraction of these cCREs are bound by CTCF in the mouse brain (Figure 5)^43^. Recently, it was shown that CTCF is involved in promoter selection in the proto-cadherin gene cluster by promoting enhancer-promoter looping^55,56^. Our data now suggest that CTCF could regulate a broader set of neuronal genes than previously demonstrated^55,76^, which need to be verified in future experiments. In addition, the RFX family of transcription factors was described to regulate cilia in sensory neurons^58^. Our data suggest a more widespread role for RFX family of transcription factors in the brain in regulation of synaptic transmission. Consistent with this proposal, deletion of *Rfx4* gene in mouse was shown to severely disrupt neural development^77^. We have previously shown that RFX motif was enriched in elements that were more accessible after birth compared to prenatal time points in both GABAergic and glutamatergic neuronal types^31^. RFX was also found to be strongly enriched in mouse forebrain enhancers with increased activity after birth^78^. Similar to CTCF and RFX, the MEF2 family transcription factors have been demonstrated to play roles in neurodevelopment and mental disorders^60^. The MEF2 motif was enriched at enhancers with higher chromatin accessibility in late forebrain development in mice coinciding with synapse formation^78^.

Thus, our results are consistent with the notion that cell identity is encoded in distal enhancer sequences, executed by sequence-specific transcription factors during different stages of brain development. The reference maps of cCREs for the mouse cerebrum would not only help to understand the mechanisms of gene regulation in different brain cell types, but also enable targeting and purification of specific neuronal or non-neuronal cell types or targeted gene therapy^28,79^. In addition, the maps of cCREs in the mouse brain would also assist the interpretation of non-coding risk variants associated with neurological diseases^73^. The datasets described here represent a rich resource for the neuroscience community to understand the molecular patterns underlying diversification of brain cell types in complementation to other molecular and anatomical data.

## ACKNOWLEDGEMENTS

We thank Josh Huang (CSHL) for critical reading of the manuscript. We thank Drs. Ramya Raviram, Yanxiao Zhang, Guoqiang Li and James Hocker for discussions and all other members of the Ren laboratory for valuable inputs. We would like to extend our gratitude to the QB3 Macrolab at UC Berkeley for purification of the Tn5 transposase. This study was supported by NIH Grant U19MH11483. Work at the Center for Epigenomics was also supported by the UC San Diego School of Medicine.

## AUTHOR CONTRIBUTIONS

Study was conceived by: B.R., M.M.B., J.R.E, Study supervision: B.R., Contribution to data generation: S.P., X.H., J.Y.H., X.W., D.G., S.K., J.L., M.M.B., Contribution to data analysis: Y.E.L., K.Z., Z.Z., R.F., Y.Q., O.P., Y.Y., H.L., E.A.M., Contribution to web portal: Y.E.L., Z.Z., B.L., Contribution to data interpretation: Y.E.L., S.P., B.R., J.R.E., M.B., E.A.M., Contribution to writing the manuscript: Y.E.L., S.P., B.R. All authors edited and approved the manuscript.

---

**Extended Data Figure 1:**
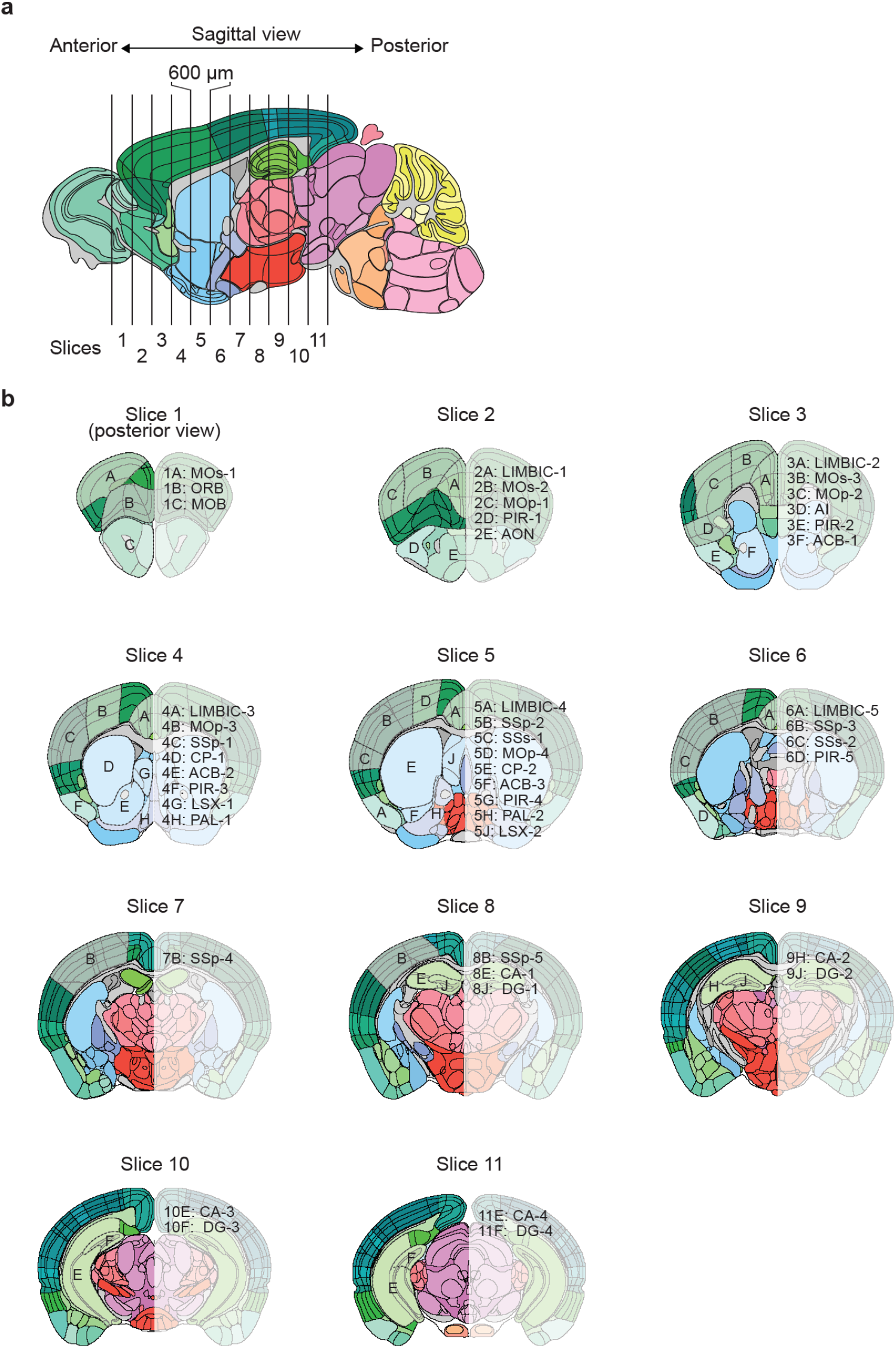
Maps of mouse brain regions that were dissected in the current study. **a** Schematic of brain sample dissection strategy. Mouse brains were cut into 600 μm thick coronal slices; **b** 45 regions were dissected from eleven coronal slices according to the Allen Brain Reference Atlas^32^. Shown is the frontal view of slice 1-11 and isolated regions. For example, dissection region 1A: MOs-1 denotes part 1 of the secondary motor cortex (MOs) region which corresponds to region A from slice 1. A detailed list of regions can be found in Supplementary Table 1.

**Extended Data Figure 2:**
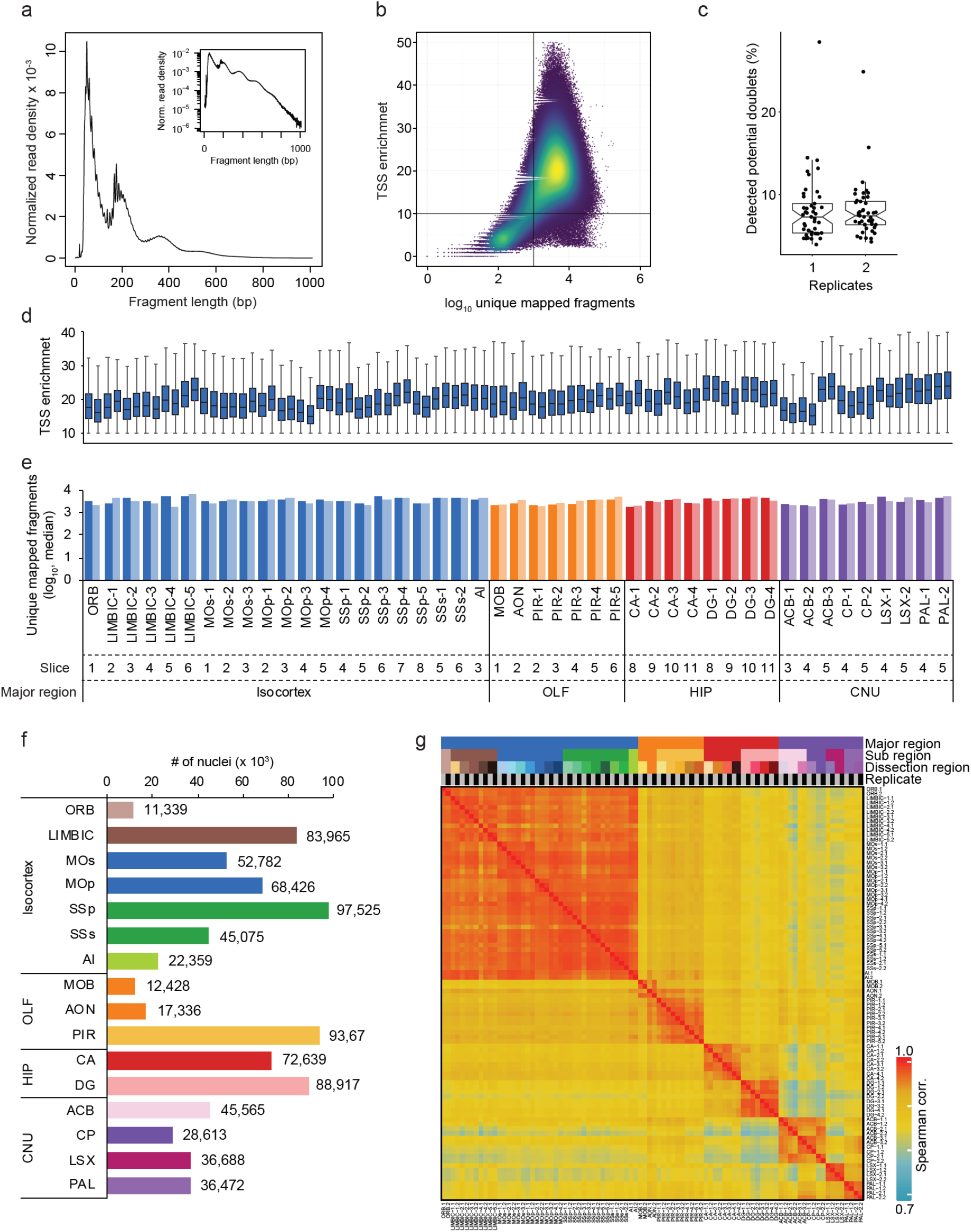
Quality metrics of snATAC-seq datasets. **a** Fragment size distribution of a typical snATAC-seq library. **b** Dot-blot illustrating fragments per nucleus and individual TSS (transcriptional start site) enrichment. Nuclei in the upper right quadrant were selected for analysis. **c** Fraction of potential barcode collisions detected in snATAC-seq libraries using a modified version of Scrublet^33^ (the box is drawn from lower quartile (Q1) to upper quartile (Q3) with a horizontal line drawn in the middle to denote the median, whiskers with maximum 1.5 IQR). Potential barcode collisions were removed for downstream processing. **d** Distribution of TSS enrichment (the box is drawn from lower quartile (Q1) to upper quartile (Q3) with a horizontal line drawn in the middle to denote the median, whiskers with maximum 1.5 IQR) and **e** number of uniquely mapped fragments/nucleus for individual libraries. **f** Number of nuclei passing quality control for sub-regions. **g** Spearman correlation matrix of snATAC-seq libraries.

**Extended Data Figure 3:**
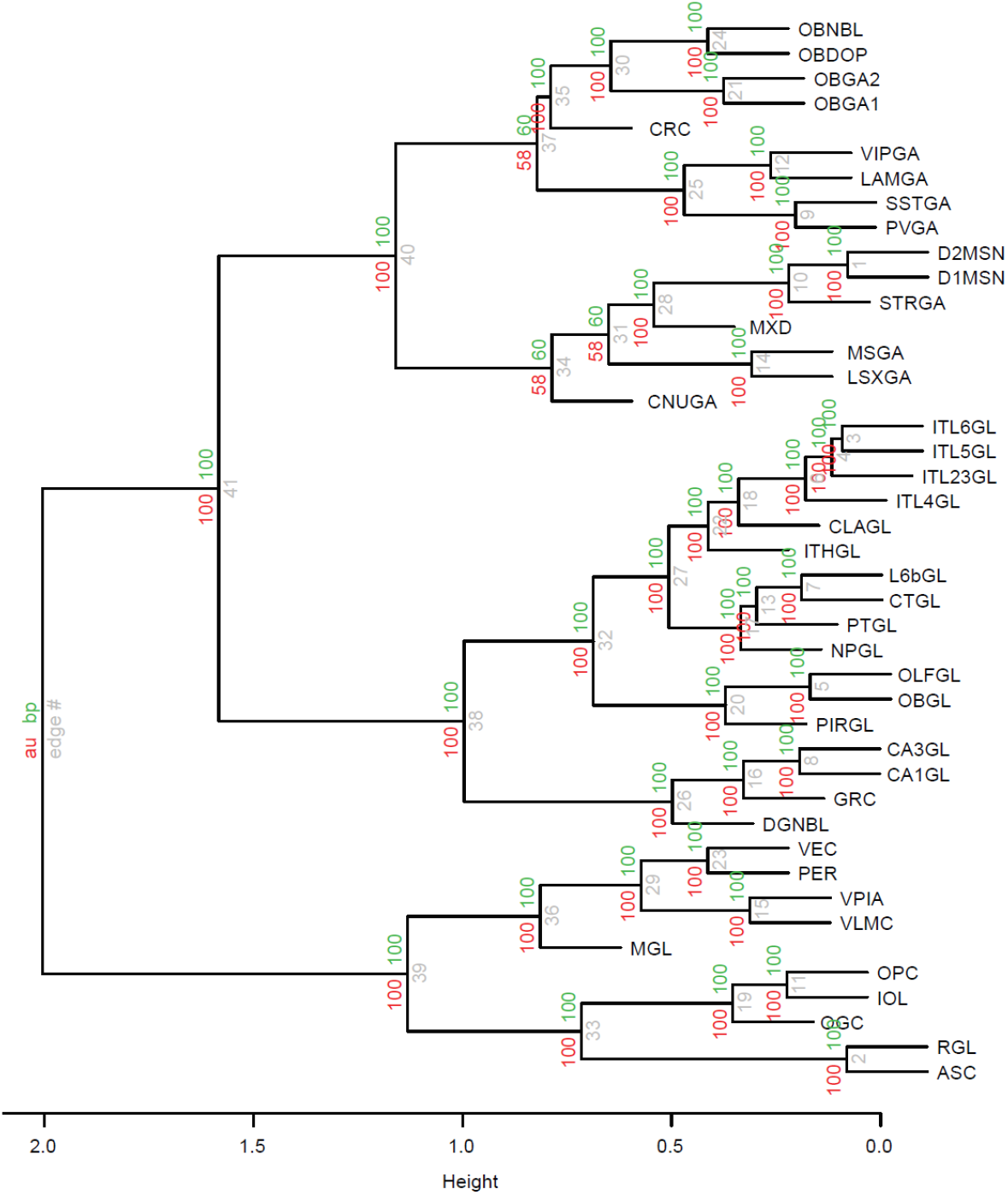
Hierarchical dendrogram of the major cell types. Dendrogram for major cell types was constructed using 1000 rounds of bootstrapping for major cell types using R package pvclust^83^. Nodes are labeled in grey, approximately unbiased (AU) p-values (in red) and bootstrap probability (BP) values (in green) are labeled at the shoulder of the nodes, respectively. For a full list and description of cell cluster labels see Supplementary Table 3.

**Extended Data Figure 4:**
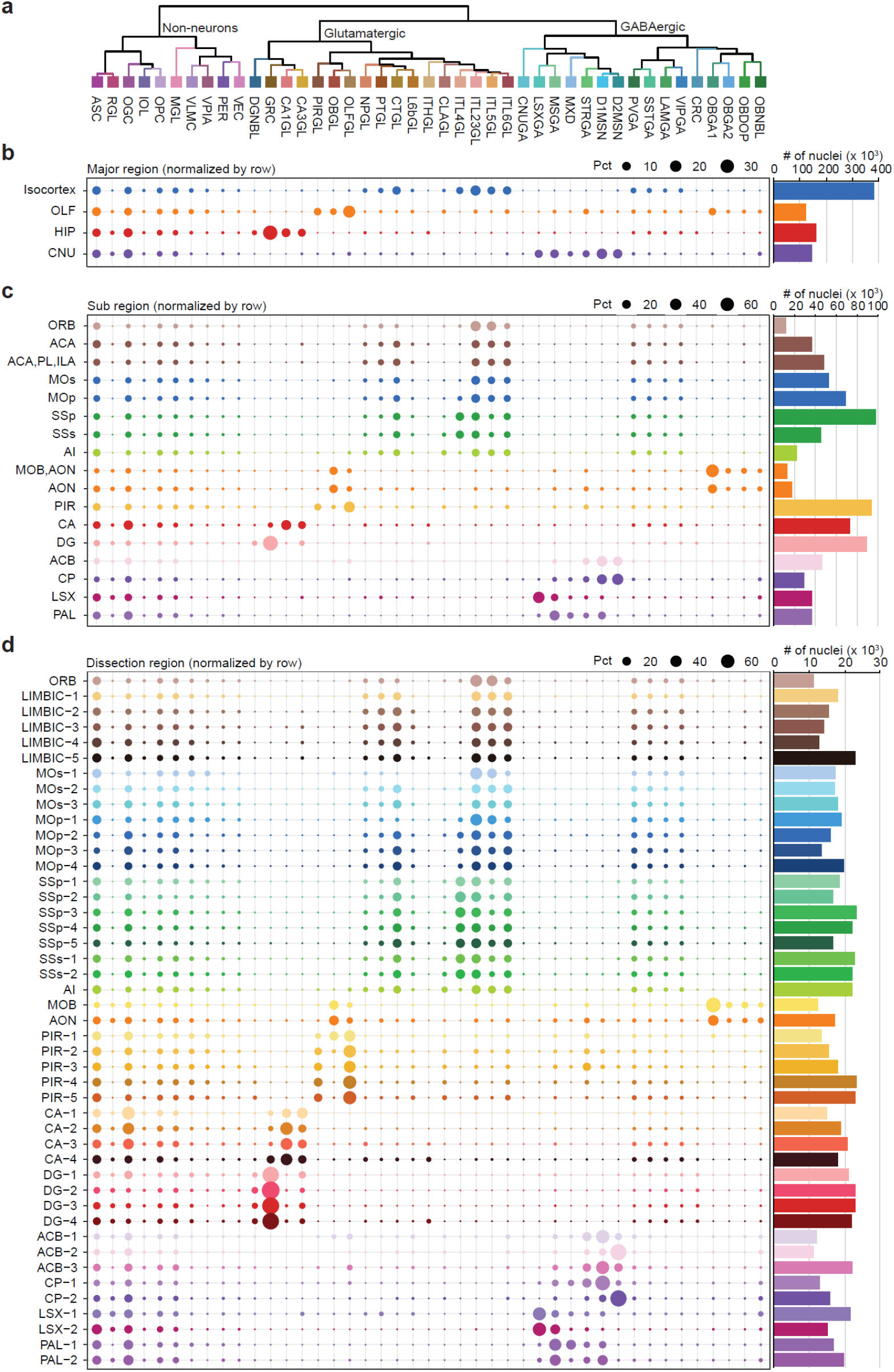
Relative cell cluster proportion at region resolution. **a** Cluster dendrogram based on chromatin accessibility. **b-d** Major cell-type composition in **b** the four major regions, **c** the sub-regions and **d** the dissected regions. Indicated are row normalized percentages (pct) of clusters per major region and the total number of nuclei for each major region. Bar plots to the right show total number of nuclei sampled for each region.

**Extended Data Figure 5:**
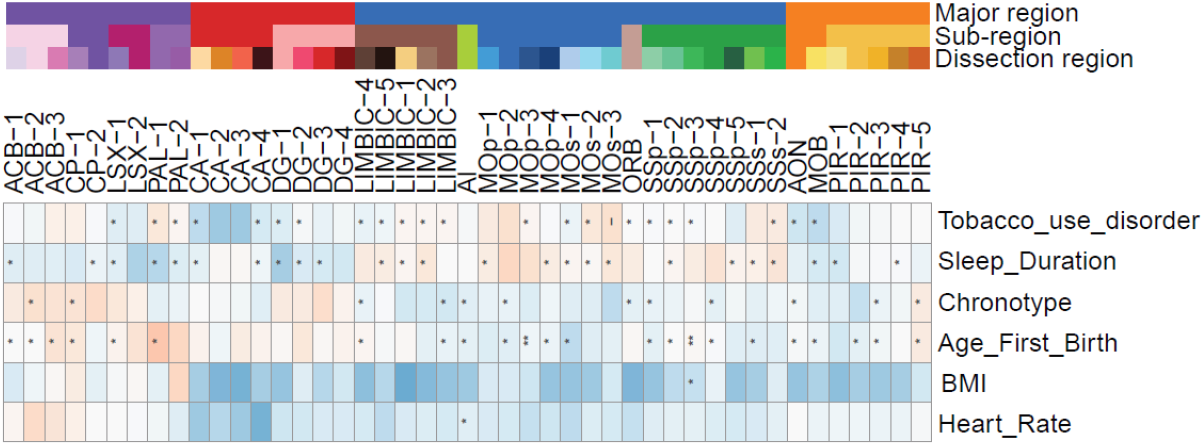
GWAS enrichment for additional traits in open chromatin of distinct cell types. Brain region specific enrichment of indicated GWAS traits (* FDR < 0.05, ** FDR <0.01, ***FDR < 0.001). Displayed are all brain regions and all tested phenotypes with at least one significant association.

## TABLES

**Supplementary Table1:** Sample and dissection summary

**Supplementary Table 2:** Metadata table for nuclei

**Supplementary Table 3:** Cell cluster annotation

**Supplementary Table 4:** Overlap score for integration of snATAC-seq and scRNA-seq clusters

**Supplementary Table 5:** List of the genomic locations of cCREs

**Supplementary Table 6:** Cluster assignment of cCREs

**Supplementary Table 7:** Association of *cis* regulatory modules with major cell types

**Supplementary Table 8:** Module assignment of cCREs

**Supplementary Table 9:** Known motif enrichment in *cis* regulatory modules

**Supplementary Table 10:** Summary of gene-cCRE correlations

**Supplementary Table 11:** Association of modules with joint cell clusters

**Supplementary Table 12:** Association of modules with individual putative enhancers

**Supplementary Table 13:** Gene Ontology analysis of candidate target genes of putative enhancers

**Supplementary Table 14:** Known motif enrichment in putative enhancers

**Supplementary Table 15:** *De novo* motif enrichment in module M1 putative enhancers

**Supplementary Table 16:** Gene Ontology analysis of candidate target gene of putative enhancers with motif sites in module M1

**Supplementary Table 17:** Known motif enrichment in candidate target promoters of putative enhancers

**Supplementary Table 18:** Primer sequences and nuclei barcodes for version 1 and 2 indexing schemes.

## METHODS

### Tissue preparation and nuclei isolation

All experimental procedures using live animals were approved by the SALK Institute Animal Care and Use Committee under protocol number 18-00006. Adult C57BL/6J male mice were purchased from Jackson Laboratories. Brains were extracted from 56-63 day old mice and sectioned into 600 μm coronal sections along the anterior-posterior axis in ice-cold dissection media.^14,15^ Specific brain regions were dissected according to the Allen Brain Reference Atlas^32^ (Extended Data Figure 1) and nuclei isolated as described.^15^

### Single nucleus ATAC-seq

Single nucleus ATAC-seq was performed as described with steps optimized for automation^15,31,34^. A step-by-step-protocols for library preparation is available here: https://www.protocols.io/view/sequencing-open-chromatin-of-single-cell-nuclei-sn-pjudknw/abstract.

Brain nuclei were pelleted with a swinging bucket centrifuge (500 × g, 5 min, 4°C; 5920R, Eppendorf). Nuclei pellets were resuspended in 1 ml nuclei permeabilization buffer (5 % BSA, 0.2 % IGEPAL-CA630, 1mM DTT and cOmpleteTM, EDTA-free protease inhibitor cocktail (Roche) in PBS) and pelleted again (500 × g, 5 min, 4°C; 5920R, Eppendorf). Nuclei were resuspended in 500 μL high salt tagmentation buffer (36.3 mM Tris-acetate (pH = 7.8), 72.6 mM potassium-acetate, 11 mM Mg-acetate, 17.6% DMF) and counted using a hemocytometer. Concentration was adjusted to 1,000-4,500 nuclei/9 μl, and 1,000-4,500 nuclei were dispensed into each well of a 96-well plate. For tagmentation, 1 μL barcoded Tn5 transposomes^34^ were added using a BenchSmart™ 96 (Mettler Toledo, RRID:SCR_018093, Supplementary Table 18), mixed five times and incubated for 60 min at 37 °C with shaking (500 rpm). To inhibit the Tn5 reaction, 10 μL of 40 mM EDTA were added to each well with a BenchSmart™ 96 (Mettler Toledo, RRID:SCR_018093) and the plate was incubated at 37 °C for 15 min with shaking (500 rpm). Next, 20 μL 2 × sort buffer (2 % BSA, 2 mM EDTA in PBS) were added using a BenchSmart™ 96 (Mettler Toledo, RRID:SCR_018093). All wells were combined into a FACS tube and stained with 3 μM Draq7 (Cell Signaling). Using a SH800 (Sony), 20 nuclei were sorted per well into eight 96-well plates (total of 768 wells) containing 10.5 μL EB (25 pmol primer i7, 25 pmol primer i5, 200 ng BSA (Sigma). Preparation of sort plates and all downstream pipetting steps were performed on a Biomek i7 Automated Workstation (Beckman Coulter, RRID:SCR_018094). After addition of 1 μL 0.2% SDS, samples were incubated at 55 °C for 7 min with shaking (500 rpm). 1 μL 12.5% Triton-X was added to each well to quench the SDS. Next, 12.5 μL NEBNext High-Fidelity 2× PCR Master Mix (NEB) were added and samples were PCR-amplified (72 °C 5 min, 98 °C 30 s, (98 °C 10 s, 63 °C 30 s, 72°C 60 s) × 12 cycles, held at 12 °C). After PCR, all wells were combined. Libraries were purified according to the MinElute PCR Purification Kit manual (Qiagen) using a vacuum manifold (QIAvac 24 plus, Qiagen) and size selection was performed with SPRI Beads (Beckmann Coulter, 0.55x and 1.5x). Libraries were purified one more time with SPRI Beads (Beckmann Coulter, 1.5x). Libraries were quantified using a Qubit fluorimeter (Life technologies, RRID:SCR_018095) and the nucleosomal pattern was verified using a Tapestation (High Sensitivity D1000, Agilent). Libraries generated with indexing version 1^34^ (Supplementary Table 1) were sequenced on a HiSeq2500 sequencer (RRID:SCR_016383, Illumina) using custom sequencing primers, 25% spike-in library and following read lengths: 50 + 43 + 37 + 50 (Read1 + Index1 + Index2 + Read2). Libraries generated with indexing version 2 (Supplementary Table 1) were sequenced on a HiSeq4000 (RRID:SCR_016386, Illumina) using custom sequencing primers with following read lengths: 50 + 10 + 12 + 50 (Read1 + Index1 + Index2 + Read2). Indexing primers and sequencing primers are in Supplementary Table 18. The nuclei indexing version (v1 or v2) used for each library is indicated in Supplementary Table 1.

### Nuclei indexing schemes

To generate snATAC-seq libraries we used initially an indexing scheme as described before (Version 1).^29,31^ Here, 16 p5 and 24 p7 indexes were combined to generate an array of 384 indexes for tagmentation and 16 i5 as well as 48 i7 indexes were combined for an array of 768 PCR indexes. Due to this library design, it is required to sequence all four indexes to assign a read to a specific nucleus with long reads and a constant base sequence for both indices reads between i and p barcodes. Therefore, the resulting libraries were sequenced with 25% spike-in library on a HiSeq2500 (RRID:SCR_016383) and these read lengths: 50+43+37+50.^31^

To generate libraries compatible with other sequencers and not requiring spike-in libraries or custom sequencing recipes, we modified the library scheme (Version 2). For this, we used 384 individual indices for T7 and combined with one T5 with a universal index sequence for tagmentation (for a total of 384 tagmentation indexes). For PCR, we used 768 different i5 indexes and combined with a universal i7 primer index sequence. Tagmentation indexes were 10 bp and PCR indexes 12 bp long. We made sure, that the hamming distance between every two barcodes was >=4, the GC content between 37.5-62.5 % and the number of repeats <= 3. The resulting libraries were sequenced on a HiSeq4000 with custom primers and these read lengths: 50+10+12+50 (Supplementary Table 18).

### Processing and alignment of sequencing reads

Paired-end sequencing reads were demultiplexed and the cell index transferred to the read name. Sequencing reads were aligned to mm10 reference genome using bwa^84^. After alignment, we used the R package ATACseqQC (1.10.2)^85^ to check for fragment length contribution which is characteristic for ATAC-seq libraries. Next, we combined the sequencing reads to fragments and for each fragment we performed following quality control: 1) Keep only fragments quality score MAPQ > 30; 2) Keep only the properly paired fragments with length <1000bp. 3) PCR duplicates were further removed with SnapTools (https://github.com/r3fang/SnapTools, RRID:SCR_018097)^34^. Reads were sorted based on the cell barcode in the read name.

### TSS enrichment calculation

Enrichment of ATAC-seq accessibility at TSSs was used to quantify data quality without the need for a defined peak set. The method for calculating enrichment at TSS was adapted from previously described. TSS positions were obtained from the GENCODE database v16 (RRID:SCR_014966)^40^. Briefly, Tn5 corrected insertions (reads aligned to the positive strand were shifted +4 bp and reads aligned to the negative strand were shifted −5 bp) were aggregated ±2,000 bp relative (TSS strand-corrected) to each unique TSS genome wide. Then this profile was normalized to the mean accessibility ±1,900-2,000 bp from the TSS and smoothed every 11bp. The max of the smoothed profile was taken as the TSS enrichment.

### Doublet removal

We used a modified version of Scrublet (RRID:SCR_018098)^33^ to remove potential doublets for every dataset independently. Peaks were called using MACS2 for aggregate accessibility profiles on each sample. Next, cell-by-peak count matrices were calculated and used as input, with default parameters. Doublet scores were calculated for both observed nuclei {x_j_} and simulated doublets {y_j_} using Scrublet (RRID:SCR_018098)^33^. Next, a threshold θ is selected based on the distribution of {y_j_}, and observed nuclei with doublet score larger than θ are predicted as doublets. To determine θ, we fit a two-component mixture distribution by using function normalmixEM from R package mixtools. The lower component contained majority of embedded doublet types, and the other component contained majority of neo-typic doublets (collision between nuclei from different clusters. We selected the threshold θ where the *p_A_ -pdf*(*x*, μ_1_ σ_1_) = *p*_2_ · *pdf*(*x*, μ_2_, σ_2_). This value suggested that the nuclei have same chance of belonging to both classes.

### Clustering and cluster annotation

We used an iterative clustering strategy using the snapATAC package (RRID:SCR_018097) with slight modifications as detailed below.^34^ For round 1 clustering, we clustered and finally merged single nuclei to three main cell classes: non-neurons, GABAergic neurons and glutamatergic neurons. For each main cell class, we preformed another round of clustering to identify major cell types. Last, for each major cell types, we performed a third round of clustering to find sub-types.

Detailed description for every step is listed below:

1. Nuclei filtering Nuclei with >=1,000 uniquely mapped fragments and TSS (transcription start site) enrichment >10 were filtered for individual dataset. Second, potential barcode collisions were also removed for individual dataset.
2. Feature bin selection First, we calculated a cell-by-bin matrix at 500 kb resolution for every dataset independently and subsequently merged the matrices. Second, we converted the cell-by-bin count matrix to a binary matrix. Third, we filtered out any bins overlapping with the ENCODE blacklist (mm10, http://mitra.stanford.edu/kundaje/akundaje/release/blacklists/mm10-mouse/mm10.blacklist.bed.gz). Fourth, we focused on bins on chromosomes 1-19, X and Y. Last, we removed the top 5% bins with the highest read coverage from the count matrix.
3. Dimensionality reduction SnapATAC applies a nonlinear dimensionality reduction method called diffusion maps, which is highly robust to noise and perturbation.^34^ However, the computational time of the diffusion maps algorithm scales exponentially with the increase of number of cells. To overcome this limitation, we combined the Nyström method (a sampling technique)^86^ and diffusion maps to present Nyström Landmark diffusion map to generate the lowdimensional embedding for large-scale dataset. A Nyström landmark diffusion maps algorithm includes three major steps:

1. sampling: sample a subset of K (K≪ N) cells from N total cells as “landmarks”.
2. embedding: compute a diffusion map embedding for K landmarks;
3. extension: project the remaining N-K cells onto the low-dimensional embedding as learned from the landmarks to create a joint embedding space for all cells. Having more than 800,000 single nuclei at the beginning, we decided to apply this strategy on the level 1 and 2 clustering. 10,000 cells were sampled as landmarks and the remaining query cells were projected onto the diffusion maps embedding of landmarks. Later for the level III clustering, diffusion map embeddings were directly calculated from all nuclei.
4. Principal Component (PC) selection To determine the number of principal components to include for downstream analysis, we generated “Elbow plot”, to rank all principal components based on the percentage of variance explained by each one. For each round of clustering, we selected the top 10-20 principal components that captured the majority of the variance.
5. Graph-based clustering Using the selected significant components, we next construct a K Nearest Neighbor (KNN) Graph. Each cell is a node and the k-nearest neighbours of each cell were identified according to the Euclidian distance and edges were drawn between neighbours in the graph. Next, we applied the Leiden algorithm on the KNN graph using python package leidenalg (https://github.com/vtraag/leidenalg)^87^. We tested different ‘resolution_parameter’ parameters (step between 0 and 1 by 0.1) to determine the optimal resolution for different cell populations. For each resolution value, we tested if there was clear separation between nuclei. To do so, we generated a cell-by-cell consensus matrix in which each element represents the fraction of observations two nuclei are part of the same cluster. A perfectly stable matrix would consist entirely of zeros and ones, meaning that two nuclei either cluster together or not in every iteration. The relative stability of the consensus matrices can be used to infer the optimal resolution. To this end, we generated a consensus matrix based on 300 rounds of Leiden clustering with randomized starting seed *s*. let M^s^ denote the N × N connectivity matrix resulting from applying Leiden algorithm to the dataset D^s^ with different seeds. The entries of M^s^ are defined as follows:

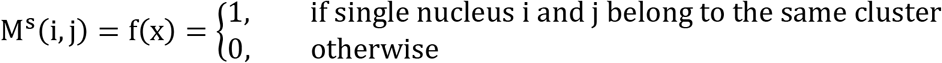 Let I^s^ be the N × N identicator matrix where the (i, j)-th entry is equal to 1 if nucleus i and j are in the same perturbed dataset D^s^, and 0 otherwise. Then, the consensus matrix C is defined as the normalised sum of all connectivity matrices of all the perturbed D^s^.

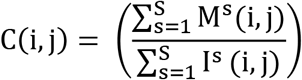 The entry (i, j) in the consensus matrix is the number of times single nucleus i and j were clustered together divided by the total number of times they were selected together. The matrix is symmetric, and each element is defined within the range [0,1]. We examined the cumulative distribution function (CDF) curve and calculated proportion of ambiguous clustering (PAC) score to quantify stability at each resolution. The resolution with a local minimal of the PAC scores denotes the parameters for the optimal clusters. In the case these were multiple local minimal PACs, we picked the one with higher resolution. Finally, for every cluster, we tested whether we could identify differential features compared to all other nuclei (background) and to the nearest nuclei (local background) using the function ‘findDAR’.
6. Visualization For visualization we applied Uniform Manifold Approximation and Projection (UMAP)^80^.

### Regional specificity

For each cell type, fraction of nuclei is first calculated from each brain regions. Then, we use function ‘entropyDiversity’ from R package BioQC (cite) to calculate regional diversity for each cell types and minus the value by 1 as specificity.

### Identification of reproducible peak sets in each cell cluster

We performed peak calling according to the ENCODE ATAC-seq pipeline (https://www.encodeproject.org/atac-seq/). For every cell cluster, we combined all properly paired reads to generate a pseudobulk ATAC-seq dataset for individual biological replicates. In addition, we generated two pseudo-replicates which comprise half of the reads from each biological replicate. We called peak for each of the four dataset and a pool of both replicates independently. Peak calling was performed on the Tn5-corrected single-base insertions using the MACS2^39^ with these parameters: --shift −75 -- extsize 150 --nomodel --call-summits --SPMR --keep-dup all -q 0.01. Finally, we extended peak summits by 250 bp on either side to a final width of 501 bp for merging and downstream analysis. To generate a list of reproducible peaks, we kept peaks that 1) were detected in the pooled dataset and overlapped >=50% of peak length with a peak in both individual replicates or 2) were detected in the pooled dataset and overlapped >=50% of peak length with a peak in both pseudo-replicates.

To account for differences in performance of MACS2^39^ based on read depth and/or number of nuclei in individual clusters, we converted MACS2 peak scores (-log10(q-value)) to “score per million”^88^. We filtered reproducible peaks by choosing a “score per million” cut-off of 2 was used to filter reproducible peaks.

We only kept reproducible peaks on chromosome 1-19 and both sex chromosomes, and filtered ENCODE mm10 blacklist regions (mm10, http://mitra.stanford.edu/kundaje/akundaje/release/blacklists/mm10-mouse/mm10.blacklist.bed.gz). A union peak list for the whole dataset obtained by merging peak sets from all cell clusters using BEDtools (RRID:SCR_006646)^89^.

Lastly, since snATAC-seq data are very sparse, we selected only elements that were identified as open chromatin in a significant fraction of the cells in each cluster. To this end, we first randomly selected same number of non-DHS regions (~ 670k elements) from the genome as background and calculated the fraction of nuclei for each cell type that that showed a signal at these sites. Next, we fitted a zero-inflated beta model and empirically identified a significance threshold of FDR < 0.01 to filter potential false positive peaks. Peak regions with FDR < 0.01 in at least one of the clusters were included into downstream analysis.

### Computing chromatin accessibility scores

Accessibility of cCREs in individual clusters was quantified by counting the fragments in individual clusters normalized by read depth (counts per million: CPM).

For each gene, we summed counts within the gene body + 2kb upstream to calculate “gene activity score (GAS)” using Seurat (https://satijalab.org/seurat/v3.1/atacseq_integration_vignette.html, RRID:SCR_016341)^38^, GAS were used for visualization and integrative analysis with single cell RNA-seq.

### Integrative analysis of single nucleus ATAC-seq and single cell RNA-seq for mouse brain

For integrative analysis, we downloaded level 5 clustering data from the Mouse Brain Atlas website (http://mousebrain.org)^1^. First, we filtered brain regions that matched samples profiled in this study using these attributes for “Region”: “CNS”, “Cortex”, “Hippocampus”, “Hippocampus,Cortex”, “Olfactory bulb”, “Striatum dorsal”, “Striatum ventral”, “Dentate gyrus”, “Striatum dorsal,Striatum ventral”, “Striatum dorsal, Striatum ventral, Dentate gyrus”, “Pallidum”, “Striatum dorsal, Striatum ventral, Amygdala”, “Striatum dorsal, Striatum ventral”, “Telencephalon”, “Brain”, “Sub ventricular zone, Dentate gyrus” Second, we manually subset cell types into three groups by checking the attribute in “Taxonomy_group”: Non-neurons: “Vascular and leptomeningeal cells”, “Astrocytes”, “Oligodendrocytes”, “Ependymal cells”, “Microglia”, “Oligodendrocyte precursor cells”, “Olfactory ensheathing cells”, “Pericytes”, “Vascular smooth muscle cells”, “Perivascular macrophages”, “Dentate gyrus radial glia-like cells”, “Subventricular zone radial glia-like cells”, “Vascular smooth muscle cells”, “Vascular endothelial cells”, “Vascular and leptomeningeal cells”; GABAergic neurons: “Non-glutamatergic neuroblasts”, “Telencephalon projecting inhibitory neurons”, “Olfactory inhibitory neurons”, “Glutamatergic neuroblasts”, “Cholinergic and monoaminergic neurons”, “Di- and mesencephalon inhibitory neurons”, “Telencephalon inhibitory interneurons”, “Peptidergic neurons”; Glutamatergic neurons: “Dentate gyrus granule neurons”, “Di- and mesencephalon excitatory neurons”, “Telencephalon projecting excitatory neurons” We performed integrative analysis with single cell RNA-seq using Seurat 3.0 (RRID:SCR_016341) to compare cell annotation between different modalities^38^. We randomly selected 200 nuclei (and used all nuclei for cell cluster with <200 nuclei) from each cell cluster for integrative analysis. We first generated a Seurat object in R by using previously calculated gene activity scores, diffusion map embeddings and cell cluster labels from snATAC-seq. Then, variable genes were identified from scRNA-seq and used for identifying anchors between these two modalities. Finally, to visualize all the cells together, we co-embedded the scRNA-seq and snATAC-seq profiles in the same low dimensional space.

To quantify the similarity between cell clusters from two modalities, we calculated an overlapping score as the sum of the minimum proportion of cells/nuclei in each cluster that overlapped within each co-embedding cluster^10^. Cluster overlaps varied from 0 to 1 and were visualized as a heat map with snATAC-seq clusters in rows and scRNA-seq clusters in columns.

### Identification of *cis* regulatory modules

We used Nonnegative Matrix Factorization (NMF)^90^ to group cCREs into *cis* regulatory modules based on their relative accessibility across major clusters. We adapted NMF (Python package: sklearn^91^) to decompose the cell-by-cCRE matrix V (N×M, N rows: cCRE, M columns: cell clusters) into a coefficient matrix H (R×M, R rows: number of modules) and a basis matrix W (N×R), with a given rank R:

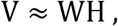

The basis matrix defines module related accessible cCREs, and the coefficient matrix defines the cell cluster components and their weights in each module. The key issue to decompose the occupancy profile matrix was to find a reasonable value for the rank R (i.e., the number of modules). Several criteria have been proposed to decide whether a given rank R decomposes the occupancy profile matrix into meaningful clusters. Here we applied two measurements “Sparseness”^92^ and “Entropy”^93^ to evaluate the clustering result. Average values were calculated from 100 times for NMF runs at each given rank with random seed, which will ensure the measurements are stable.

Next, we used the coefficient matrix to associate modules with distinct cell clusters. In the coefficient matrix, each row represents a module and each column represents a cell cluster. The values in the matrix indicate the weights of clusters in their corresponding module. The coefficient matrix was then scaled by column (cluster) from 0 to 1. Subsequently, we used a coefficient > 0.1 (~95th percentile of the whole matrix) as threshold to associate a cluster with a module.

In addition, we associated each module with accessible elements using the basis matrix. For each element and each module, we derived a basis coefficient score, which represents the accessible signal contributed by all cluster in the defined module. In addition, we also implemented and calculated a basis-specificity score called “feature score” for each accessible element using the “kim” method^93^. The feature score ranges from 0 to 1. A high feature score means that a distinct element is specifically associated with a specific module. Only features that fulfil both following criteria were retained as module specific elements:

1. feature score greater than median + 3 standard deviation;
2. the maximum contribution to a basis component is greater than the median of all contributions (i.e. of all elements of W).

### Dendrogram construction for mouse brain cell types

First, we calculated for cCRE the median accessibility per cluster and used this value as cluster centroid. Next, we calculated the coefficient of variant (CV) for the cluster centroid of each element across major cell types. Finally, we only kept variable elements with CV larger than 1.5 for dendrogram construction.

We used the set of variable features defined above to calculate a correlation-based distance matrix. Next, we performed linkage hierarchical clustering using the R package pvclust (v.2.0)^83^ with parameters method.dist=“cor” and method.hclust=“ward.D2”. The confidence for each branch of the tree was estimated by the bootstrap resampling approach.

### Motif enrichment

We performed both *de novo* and known motif enrichment analysis using Homer (v4.11, RRID:SCR_010881)^46^. For cCREs in the consensus list, we scanned a region of ± 250 bp around the center of the element. And for proximal/promoter regions, we scanned a region of ± 1000 bp around the transcriptional start site.

### GREAT analysis

Gene ontology annotation of cCREs was performed using GREAT (version 4.0.4, RRID:SCR_005807)^94^ with default parameters. GO Biological Process was used for annotations.

### Gene ontology enrichment

We perform gene ontology enrichment analysis using R package Enrichr (RRID:SCR_001575)^82^. Gene set library “GO_Biological_Process_2018” was used with default parameters. The combined score is defined as the p-value computed using the Fisher exact test multiplied with the z-score of the deviation from the expected rank.

### Predicting enhancer-promoter interactions

First, co-accessible regions are identified for all open regions in each cell cluster (randomly selected 200 nuclei, and used all nuclei for cell cluster with <200 nuclei) separately, using Cicero^49^ with following parameters: aggregation k = 10, window size = 500 kb, distance constraint = 250 kb. In order to find an optimal co-accessibility threshold for each cluster, we generated a random shuffled cCRE-by-cell matrix as background and identified co-accessible regions from this shuffled matrix. We fitted the distribution of coaccessibility scores from random shuffled background into a normal distribution model by using R package fitdistrplus^95^. Next, we tested every co-accessibility pairs and set the cut-off at co-accessibility score with empirically defined significance threshold of FDR<0.01.

CCRE outside of ± 1 kb of transcriptional start sites (TSS) in GENCODE mm10 (v16, RRID:SCR_014966).^40^ were considered distal. Next, we assigned co-accessibility pairs to three groups: proximal-to-proximal, distal-to-distal, and distal-to-proximal. In this study, we focus only on distal-to-proximal pairs. We further used RNA expression from matched T-types to filter pairs that were linked to non-expressed genes (normalized UMI > 5).

We calculated Pearson’s correlation coefficient (PCC) between gene expression and cCRE accessibility across joint RNA-ATAC clusters to examine the relationship between co-accessibility pairs. To do so, we first aggregated all nuclei/cells from scRNA-seq and snATAC-seq for every joint cluster to calculate accessibility scores (log_2_ CPM) and relative expression levels (log_2_ normalized UMI). Then, PCC was calculated for every gene-cCRE pair within a 1 Mbp window centered on the TSS for every gene. We also generated a set of background pairs by randomly selecting regions from different chromosomes and shuffling of cluster labels. Finally, we fit a normal distribution model and defined a cut-off at PCC score with empirically defined significance threshold of FDR<0.01, in order to select significant positively correlated cCRE-gene pairs.

### GWAS enrichment

To enable comparison to GWAS of human phenotypes, we used liftOver with settings “-minMatch=0.5” to convert accessible elements from mm10 to hg19 genomic coordinates.^69^ Next, we reciprocal lifted the elements back to mm10 and only kept the regions that mapped to original loci. We further removed converted regions with length > 1kb.

We obtained GWAS summary statistics for quantitative traits related to neurological disease and control traits: Heart Failure^96^, Type 1 Diabetes^97^, Age First Birth and Number Children Born^98^, Lupus^99^, Primary Biliary Cirrhosis^100^, Tiredness^101^, Crohns_Disease^102^, Inflammatory Bowel Disease^102^, Ulcerative_Colitis^102^, Asthma^103^, Attention Deficit Hyperactivity Disorder^104^, Heart Rate^105^, Celiacs Disease^106^, HOMA-B^107^, HOMA-IR^107^, Childhood Aggression^108^, Atopic Dermatitis^109^, Allergy^110^, HDL_Cholesterol^111^, LDL_Cholesterol^111^, Total Cholesterol^111^, Triglycerides^111^, Autism Spectrum Disorder^112^, Birth Weight^113^, Bipolar Disorder^114^, Multiple Sclerosis^115^, Insomnia^116^, Vitamin D^117^, Primary Sclerosing Cholangitis^118^, Vitiligo^119^, Chronotype^120^, Sleep Duration^120^, Alzheimer’s Disease^121^, BMI^122^, Neuroticism^123^, Type 2 Diabetes^124^, Stroke^125^, Fasting Glucose^126^, Fasting Insulin^126^, Child Sleep Duration^127^, Coronary Artery Disease^128^, Atrial Fibrillation^129^, Rheumatoid Arthritis^130^, Educational Attainment^131^, Chronic Kidney Disease^132^, Obsessive Compulsive Disorder^133^, Post Traumatic Stress Disorder^134^, Schizophrenia^135^, Age At Menopause^136^, Age At Menarche^137^, Tobacco use disorder (ftp://share.sph.umich.edu/UKBB_SAIGE_HRC/, Phenotype code: 318)^138^, Intelligence^139^, Alcohol Usage^140^, Fasting Proinsulin^141^, Head Circumference^142^, Microalbuminuria^143^, Extraversion^144^, Birth Length^145^, Amyotrophic Lateral Sclerosis^146^, Anorexia Nervosa^147^, HbA1c^148^, Major Depressive Disorder^149^, Height^150^.

We prepared summary statistics to the standard format for Linkage disequilibrium (LD) score regression. We used homologous sequences for each major cell types as a binary annotation, and the superset of all candidate regulatory peaks as the background control. For each trait, we used cell type specific (CTS) LD score regression (https://github.com/bulik/ldsc) to estimate the enrichment coefficient of each annotation jointly with the background control^70^.

### External datasets

We listed all the datasets we used in this study for intersection analysis: rDHS regions for both hg19 and mm10 are obtained from SCREEN database (https://screen.encodeproject.org)^41,42^.

ChromHMM^43,45^ states for mouse brain are download from GitHub (https://github.com/gireeshkbogu/chromatin_states_chromHMM_mm9), and coordinates are LiftOver (https://genome.ucsc.edu/cgi-bin/hgLiftOver) to mm10 with default parameters^69^.

PhastCons^81^ conserved elements were download from the UCSC Genome Browser (http://hgdownload.cse.ucsc.edu/goldenpath/mm10/phastCons60way/).

CTCF binding sites are download from Mouse Encode Project^43^ http://chromosome.sdsc.edu/mouse/). CTCF binding sites from cortex and olfactory bulb were used in this study. Peaks are extended ± 500 bp from the loci of peak summits and used LiftOver to mm10^69^.

### Statistics

No statistical methods were used to predetermine sample sizes. There was no randomization of the samples, and investigators were not blinded to the specimens being investigated. However, clustering of single nuclei based on chromatin accessibility was performed in an unbiased manner, and cell types were assigned after clustering. Low-quality nuclei and potential barcode collisions were excluded from downstream analysis as outlined above. For significance of ontology enrichments using GREAT, Bonferroni-corrected binomial p values were used^94^. For ontology enrichment using Enrichr the combined score which represents the product of the p-value computed using the Fisher exact test multiplied with the z-score of the deviation from the expected rank was used^82^. For significance testing of enrichment of *de novo* motifs, a hypergeometric test was used without correction for multiple testing^46^.

### Data availability

Demultiplexed data can be accessed via the NEMO archive (NEMO, RRID:SCR_016152) here: http://data.nemoarchive.org/biccn/grant/cemba/ecker/chromatin/scell/raw/ Processed data are available on our web portal and can be explored here: http://catlas.org/mousebrain

### Code availability

Custom code and scripts used for analysis can be accessed here: https://github.com/YoungLeeBBS/snATACutils and https://github.com/r3fang/SnapATAC.

